# Mnemonic maps of visual space in human prefrontal cortex

**DOI:** 10.1101/2025.10.17.683147

**Authors:** Zhengang Lu, Logan T. Dowdle, Kendrick N. Kay, Clayton E. Curtis

## Abstract

Neural theories of how the prefrontal cortex (PFC) supports working memory rely on evidence from decades of pioneering macaque research. In some respects, efforts to translate these animal models of working memory in human PFC using neuroimaging have largely failed. One possible explanation, before concluding key non-homologies between the species, is that previous neuroimaging studies used resolutions too coarse to be sensitive to intermixed distributions of neurons tuned to memorized features. To resolve this concern, we scanned human PFC at 900 micron resolution using 7T fMRI. We found that population activity in retinotopically-organized superior precentral sulcus–rather than the predicted midlateral PFC–persists during memory, encodes fine-grained information about memorized items, predicts behavioral errors in memory, and forms a stable subspace with a topological organization yoked to visual space. These results have important implications for both functional homologies between the species and theories of how the PFC supports working memory.

## Introduction

One of neuroscience’s great successes in trying to unravel the mysteries of the prefrontal cortex (PFC) was the discovery of persistently active neurons in the macaque *principal sulcus* during working memory retention intervals ^1,2^. Persistent activity is often highly stimulus specific ^3–5^ and largely driven by the memory for stimuli in the contralateral visual field ^4^. Critically, lesions to the macaque principal sulcus that disrupt persistent activity cause working memory deficits that are primarily localized to specific portions of the contralesional visual field ^6^. These properties suggest that the principal sulcus stores information received from the visual system to support working memory. Persistent activity, in the absence of visual input, depends on recurrent excitatory connections among neurons that are similarly tuned ^7^ and for balance, broad synchronized inhibition ^8,9^. Decades of such empirical evidence from macaque lateral PFC has formed the cornerstone of highly influential computational and circuit-level theories of working memory ^9,10^.

An enormous amount of research has attempted to translate these animal models of human working memory. Surprisingly however, such efforts have largely failed to find similar working memory effects in the human midlateral PFC. The very first human neuroimaging study that used Positron Emission Tomography (PET) to measure brain activity during working memory failed to find delay period activity in or even near the intermediate frontal sulci (intFS; ^11^, the set of tertiary sulci in the human lateral PFC that are the homolog of the macaque principal sulcus ^12–14^. Thirty years of follow-up fMRI studies, some simple and some that used sophisticated machine learning or encoding techniques, have largely confirmed the original PET study’s failure to find working memory signatures in the midlateral PFC ^15–18^; but see ^19^.

A strong implication of these failures would suggest that the anatomical substrates in PFC that support working memory have evolved differently in humans and non-human primates. However, before affirming such a possibility, it is critical to rule out alternative explanations—particularly those rooted in methodological limitations rather than true species differences. Indeed, indirect measures of neuronal activity, like fMRI based BOLD signals, differ significantly from extracellular electrical activity ^20^. One major difference involves the spatial scale of measurement–single neurons with microelectrodes to the pooled activity of over a hundred thousand neurons within a conventionally-sized 3mm^3^ fMRI voxel. Thus, before concluding key non-homologies between the species, one must rule out that the spatial resolutions of previous neuroimaging studies were too coarse to be sensitive to the intrinsic distributions of neurons tuned to memorized features.

The sampling resolution required to distinguish memoranda-specific working memory signals depends on the degree to which similarly tuned neurons cluster together. To clarify this point, consider the distribution of spatial receptive fields of a population of neurons in a cortical area (Fig 1a). On the one hand, a population of neurons with similar receptive fields forming clusters that span several millimeters would require only a coarse spatial sampling that matches the broader cluster size (e.g., conventional 3mm^3^ fMRI voxel). On the other hand, a population with small clusters of similarly tuned neurons that are otherwise highly intermixed would require fine spatial sampling in order to be sensitive to the spatial tuning of the neurons (e.g., below conventional fMRI voxel sizes).

**Fig. 1.**
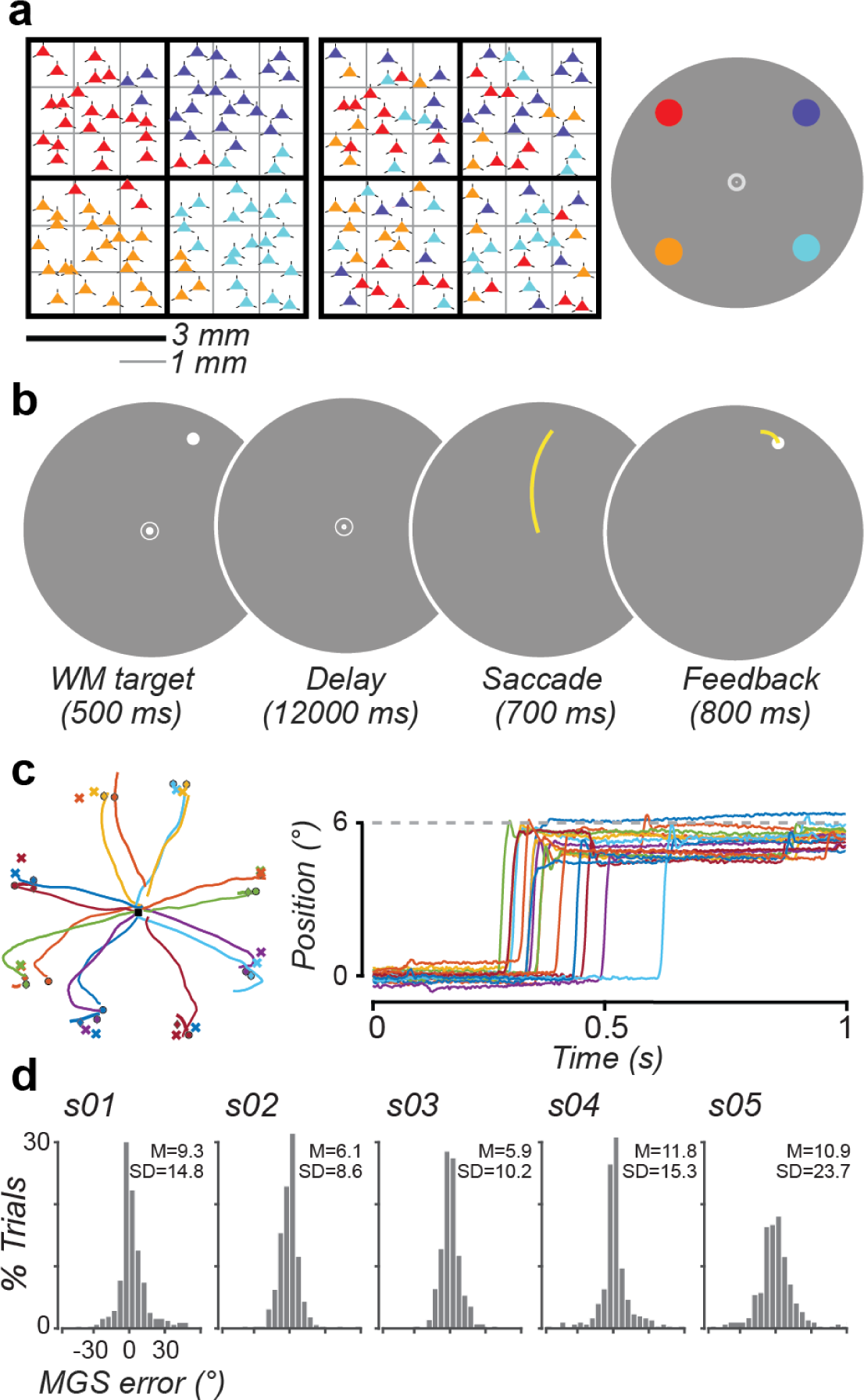
Motivation, task, and working memory behavior. **(a)** Two grids depicting the hypothetical distributions of neurons tuned to memorized locations. In the left grid, similarly tuned neurons cluster together forming patches that span several millimeters. In the right grid, they form smaller patches that only span one millimeter. Thus, detecting working memory signatures with fMRI depends on the spatial scale of sampling; coarse sampling at 3 millimeters could decode the left distribution, but fine sampling at 1 millimeter would be required to detect any item level responses from the right distribution. The colored dots in the rightmost circular display key the spatial preference of the color coded neurons (triangles). **(b)** Schematic of the memory-guided saccade task used to engage working memory. Participants maintained the location of a briefly presented target over a long retention interval and then generated a memory-guided saccade to the stored location. **(c)** Example trajectories of memory guided saccades during one scanning run (16 trials). Colored ‘x’ symbols represent actual target locations and colored dots represent saccade endpoints. **(d)** Histograms of angular component of memory errors for each participant (N=5). Insets are mean and standard deviation of memory errors (degrees) in terms of the absolute angular errors between targets and saccade endpoints. Note that memories are overall accurate, but critically show some trial-by-trial variation that may correlate with brain measures.

Several lines of evidence suggest that the resolution needed to be sensitive to selective working memory signals, if they exist in lateral PFC, is somewhere between 1 and 1.5mm, which is higher resolution than previous fMRI studies. While the median widths of cortical columns in macaque lateral PFC span 685𝜇m (range 380-1190𝜇m) ^21^, this fine of a resolution is unlikely to be required. Pairs of neurons in lateral PFC that are closer than 1-1.5mm are highly likely to have similar spatial tuning ^22,23^ and neurons with similar spatial tuning are more likely to have excitatory interconnections ^24,25^. Despite that neurons in PFC most often have strong spatial selectivity, the midlateral PFC does not appear to be topographically organized into a map of visual space ^22,26,27^ and pairs of neurons as close as 400𝜇m can have receptive fields in completely different, including the ipsilateral, parts of the visual field ^28,29^.

Here, we address these concerns regarding sampling resolution by scanning the human PFC during working memory at submillimeter (900𝜇m) voxel resolution using 7T fMRI. Using computational neuroimaging techniques, we tested whether human PFC exhibits behaviorally meaningful, hallmark signatures of working memory storage as predicted from animal studies.

## Results

Motivated by previous findings in the macaque PFC (i.e., principal sulcus), our aim was to test for similar hallmark signatures of spatial working memory in the human intFS, the homolog of macaque principal sulcus, and other lateral PFC regions. We measured brain activity associated with spatial working memory while participants performed a memory-guided saccade task that allowed direct comparisons with non-human primate neurophysiological studies ^4,6,26,28^. On each trial, participants fixated centrally while a peripheral target dot was briefly presented (500ms). After a long 12-second delay, participants executed a saccade to the memorized location of the target (Fig. 1b). We measured the angular difference between the endpoints of memory-guided saccades and the actual target positions to quantify memory performance (Fig 1c) in the scanner. Across participants, memory-guided saccades were accurate (mean absolute angular error = 8.8°; SD of absolute angular error = 14.5°; Fig 1d) and similar to previous reports ^30–32^. Importantly, there was substantial trial-to-trial variation in memory accuracy that we later will attempt to correlate with trialwise neural measures of working memory.

### Spatial selectivity in human prefrontal cortex

To identify voxels that could support spatial working memory, we first mapped spatial tuning across the frontal cortex using a population receptive field (pRF) mapping technique (See *Methods*). Characterizing spatial tuning in the frontal cortex requires cognitively demanding engagement of attention ^33^. During the pRF mapping session, participants performed a rapid serial visual presentation (RSVP) object detection task. This task required participants to detect the presence of a target object embedded in an array of objects that were rapidly changing while maintaining central fixation. The array swept across the visual field vertically and horizontally while we measured neural responses that were used to estimate the receptive fields of PFC voxels. First, this analysis revealed many voxels across lateral PFC with sensitivity to spatial positions across the visual field. Second, clusters of voxels with robust spatial selectivity were found in the superior precentral sulcus (sPCS), located in the fundus where the PCS intersects the superior frontal sulcus (Fig 2a; black outline), and the inferior precentral sulcus (iPCS), at the junction of the PCS and inferior frontal sulcus. The polar angle preferences of the voxels in both regions formed topographic maps of contralateral visual space in both hemispheres of all participants, similar to previous reports ^33–35^. In contrast to the clear and systematic maps in primary visual cortex, the maps in sPCS are coarse. The sPCS is actually composed of two maps that share a fovea ^33^ making it sometimes more difficult to discern the topography (see SupFig 1a for zoomed in views of sPCS maps). Nonetheless, we found a linear relationship within the sPCS between how close two vertices were along the cortical surface (in millimeters) and how close those two vertices were in their receptive field’s preferred polar angle (see SupFig 1b), suggesting that the area is topographically organized. While we observed voxels with spatial selectivity scattered across other portions of the lateral PFC, no other topographic maps were discernible. Specifically, no map-like topography was found in the intermediate frontal sulcus (intFS; Fig 2a, white outline). In addition to the differences in topographic organization, voxels in the sPCS when compared to intFS had stronger spatial selectivity (Fig 2b) and contained a greater proportion of spatially selective voxels (Fig 2c), especially ones with strong spatial selectivity (Fig 2d).

**Fig. 2.**
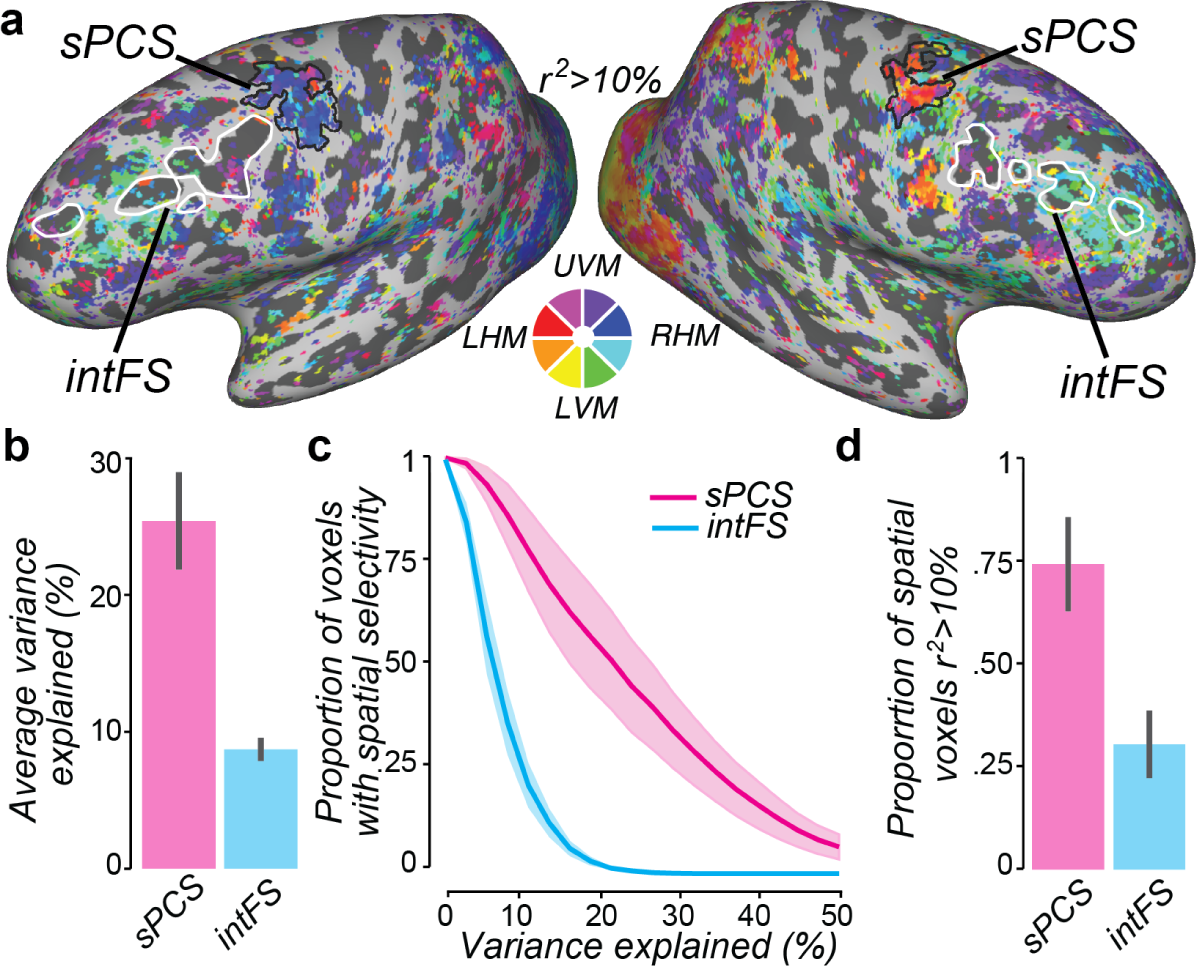
Spatial selectivity and visual field maps in human prefrontal cortex. **(a)** Using population receptive field (pRF) mapping ^33^, we modeled the receptive field properties of each cortical voxel in order to quantify the spatial selectivity of voxels and define retinotopic maps in frontal cortex for each of the participants. Retinotopic maps of contralateral visual space can be seen in superior precentral sulcus (sPCS; putative homolog of macaque frontal eye field; black outlines) for participant s01 (see SupFig 1 for other participants). The color wheel keys the polar angle of the visual field, which is overlaid on the cortical surface thresholded by variance explained (>0.1). White outlines denote anatomically defined intermediate frontal sulcus (intFS; putative homolog of macaque principal sulcus) in each hemisphere where there was no evidence of map-like structure. **(b)** Spatial selectivity of voxels in sPCS was much stronger than in intFS. Plotted is the average (± s.e.m.) amount of variance explained by the pRF model among all of the voxels in each region. **(c)** The proportion of voxels with spatial selectivity was higher in sPCS than intFS across the range of variance explained by the pRF model. **(d)** Almost 75% of voxels in the anatomically defined sPCS, compared to about 25% of voxels in intFS, had at least 10% explained variance in the pRF model.

### Persistent activity during delay period in human prefrontal cortex

Next and to identify working memory activity, we asked which parts of the human PFC exhibited sustained activity during the delay of the memory-guided saccade task. We first used general linear models (GLM) to identify voxels in PFC whose activity increased during the memory delay, while accounting for activity during the target and response epochs that bookended the delay period (see *Methods*). We observed delay period activity in PFC that was largely concentrated and most robust in the sPCS (Fig 3a). We found similarly robust delay period activity in bilateral sPCS in all participants (Sup Fig 2). Strikingly, the delay period responses overlapped precisely with the retinotopically defined maps in sPCS (Fig 3b) and over 45% of voxels in sPCS, compared to only 6% in intFS, had robust delay period activity (defined as a t-value > 4.0; Fig 3c). These data suggest a functional coupling between spatial memory and the spatially tuned voxels that form a visual field map in PFC. Supporting this coupling, we found that the magnitude of delay period activity (i.e., delay t-values) correlated with the strength of spatial tuning (i.e., pRF model variance explained) (Fig 3d, voxels pooled across participants) in the sPCS maps, r = 0.34 (p<0.001, permutation test) but weaker in the intFS maps, r = 0.08 (p<0.001, permutation test). Together, these findings indicate that the amplitude of delay period activity during WM is tightly linked to the spatial selectivity of the underlying neural populations.

**Fig. 3.**
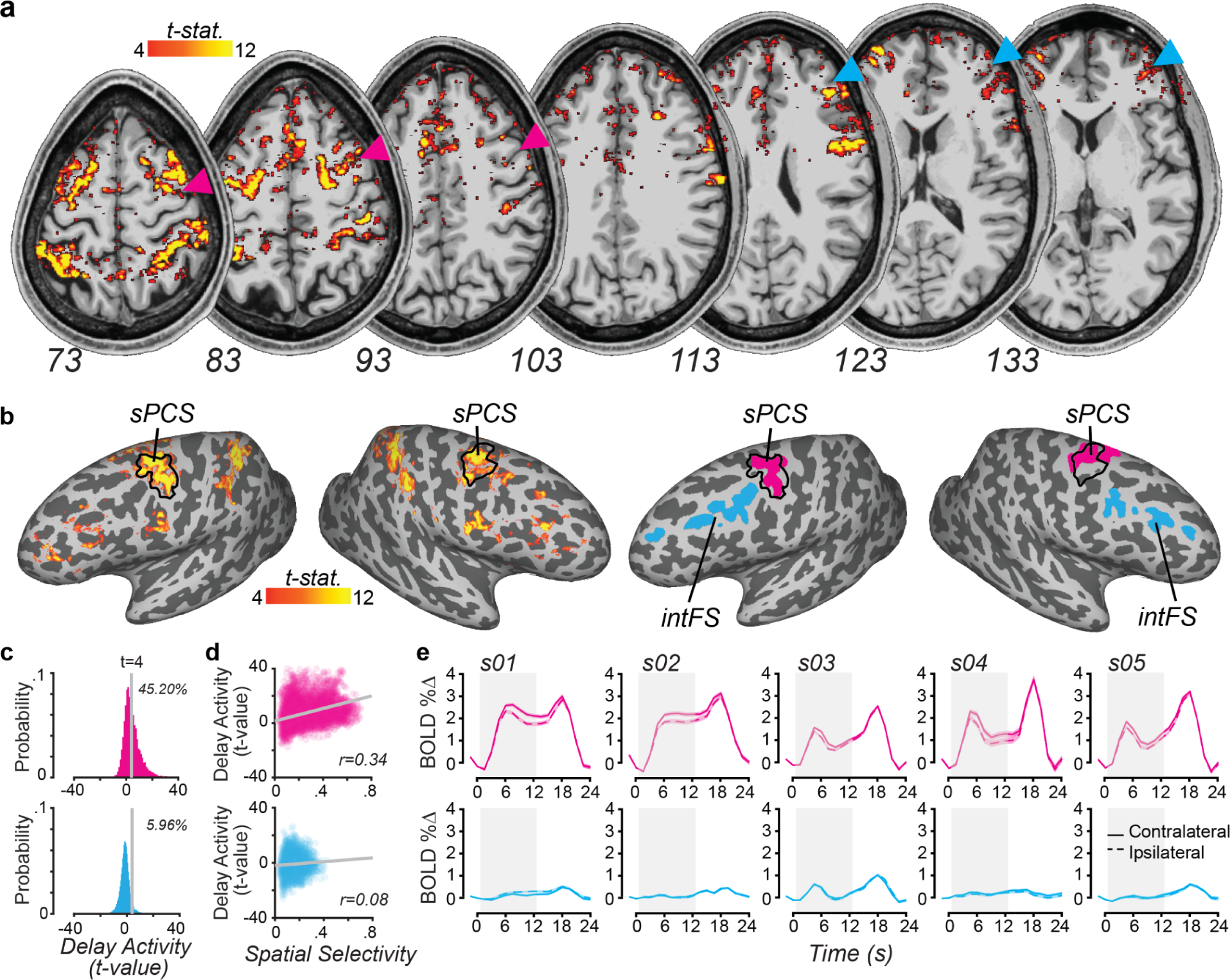
Delay period activity in human prefrontal cortex. **(a)** Statistical maps of delay period activity in an example participant (s01) overlaid on axial slices at isotropic spatial resolution of 0.9mm, thresholded at t > 4. The magenta and cyan triangles point to the superior precentral sulcus (sPCS) and intermediate frontal sulcus (intFS), respectively. **(b)** Same statistical map overlaid on the same participant’s cortical surface with Regions of interest in PFC on right. Black outlines denote retinotopically defined sPCS. Other PFC regions are in SupFig 2. The magenta and cyan colors denote the sPCS and intFS, respectively. **(c)** Probability distributions of delay period t-values. In sPCS (magenta), 45% of voxels had t-values greater than 4, compared to only 6% of voxels in intFS (cyan). Gray vertical line represents a t-value of 4. **(d)** Scatter plots showing the relationship between delay period activity (t-values) and spatial selectivity (variance explained by pRF model). In sPCS (magenta), voxels with stronger spatial selectivity had larger delay period activation. **(e)** Trial-averaged (± s.e.m.) BOLD time courses from sPCS and intFS across five participants, time-locked to trial onset. Solid and dashed lines correspond to trials with targets in the contralateral and ipsilateral visual fields, respectively. Error bars indicate s.e.m across trials. Gray shaded box denotes the delay period. Other PFC regions are in SupFig 3.

We also noted some delay period activity outside of the precentral sulcus in more anterolateral parts of the PFC. However, these responses were inconsistent across individuals, smaller in magnitude, and scattered across a small number of voxels. Notably, there were no clear and consistent delay period activations in the intFS across subjects and hemispheres (Sup Fig 2) and no correlations between the strength of delay period activity and spatial tuning like we found in sPCS.

In an attempt to be as sensitive to potential delay period activity as possible, we plotted the time-courses of BOLD activity in the top 1000 voxels with the strongest delay period activity (i.e., delay t-values; see *Methods*) in our two key PFC regions, sPCS and intFS (Fig 3b). In sPCS, a robust BOLD response time-locked to the visual target was followed by activity that persisted throughout the delay period, and followed by another robust transient response time-locked to the memory-guided saccade (Fig 3e magenta). At 7T, the amplitudes of persistent activity in sPCS were very large (i.e., 1.0-2.5% signal increase over pretrial baseline; also note very small errors bars, s.e.m across trials) compared to those previously reported at 3T (e.g., ∼0.3 - 0.5%) ^30,34,36–38^. Moreover, persistent activity in sPCS showed a clear contralateral bias, with stronger signals for targets in the contralateral visual field. In stark contrast, intFS exhibited weak or absent delay-period activity in participants, despite pre-selecting the voxels with the strongest delay period activity based on the GLM (Fig 3d blue). While the visual field map in iPCS showed robust and contralateralized delay period activity similar to sPCS, other PFC regions showed no consistent evidence for persistent delay period activity (Sup Fig 3).

### Trial-wise decoding of working memory content in prefrontal cortex

Next, we sought to understand what working memory information might be contained in the submillimeter patterns of activity in PFC that we measured with 7T fMRI and if such information relates to memory behavior. First, we used a generative encoding model combined with a Bayesian decoder (see *Methods;* ^31,39,40^) to estimate on each trial the location of the target stored in working memory. For each trial, we computed the circular mean of the decoded probability distribution for the target location and compared it to the true target location. In sPCS, decoded locations closely matched true target locations, clustering along the diagonals for all participants (Fig 4a; magenta). Subtracting the decoded locations and the true locations yielded a distribution of decoded errors, which in sPCS formed bell-shaped distributions that peaked around 0°. Decoded error distributions in sPCS differed from permutation-based null distributions (Fig 4b). In contrast, decoding from intFS produced random uniform error distributions that resembled null distributions (Fig 4a, b; cyan), indicating little if any information about the target’s location was encoded in its patterns of activity. We further quantified decoding performance using three metrics: circular correlation between decoded and true location (accuracy), mean unsigned error (precision), and standard deviation of errors (variability) (Fig 4c-e). In sPCS, decoding was accurate (average correlation r = 0.32), precise (mean unsigned error = 48.04°) and consistent (SD = 61.72°). By comparison, in intFS, decoding accuracy was low (r = 0.02), and errors were higher and more variable (mean error = 78.25°, SD = 110.30°). We also performed additional analyses to test if the working memory information might be distributed more broadly across the lateral PFC. While the decoding accuracy results indicate that sPCS contains precise information about the target locations in working memory (p = 0.0007, permutation test and Bonferroni corrected; Bayes Factor, BF₁₀= 3.42), little or no such information was found in the other ROIs (all p > 0.114), except perhaps in iPCS (p = 0.0007; BF₁₀= 1.71) and sFG (p = 0.006; BF₁₀= 1.22) (see Sup Figs 4 and 5). Previous studies reported increased delay period activity in both of these areas, but failed to decode the contents of working memory ^34,37,41^. Next, we tested if perhaps memory decoding success was not specific to sPCS, but instead depending on an anatomical gradient. Using ten bands tiling posterior-to-anterior PFC, we found that we could successfully decode the target locations only from the posterior bands which contained a large percentage of sPCS voxels (Sup Fig 6). Overall, these results demonstrate that sPCS, but not intFS or other PFC regions, robustly encodes item-level details about remembered targets on single trials.

**Fig. 4.**
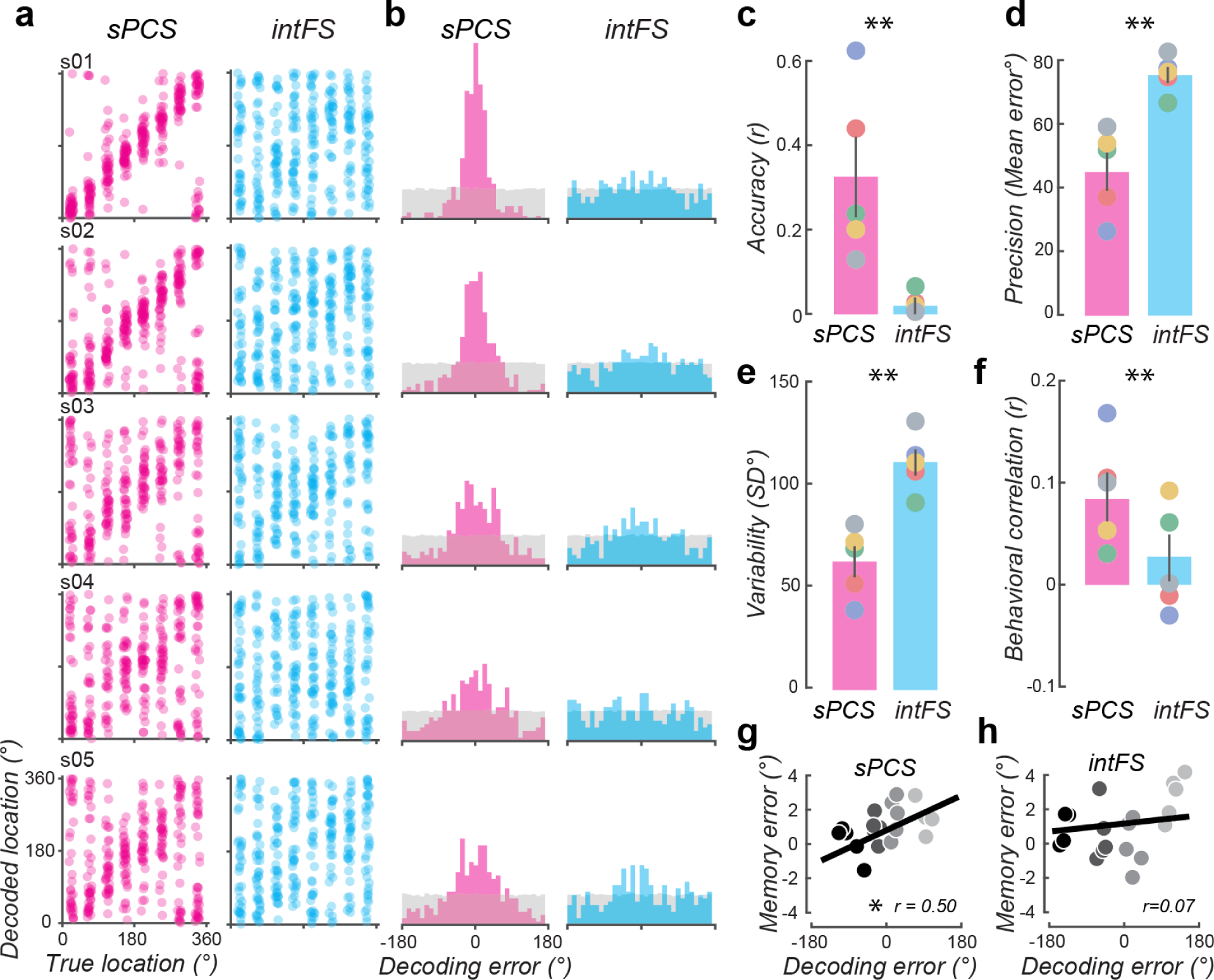
Target locations stored in working memory can be decoded from sPCS and decoding errors predict memory errors. **(a)** Decoded target locations plotted against true target locations for each participant. **(b)** Distribution of decoding errors compared to a null distribution from label-shuffled data (gray). **(c-e)** Mean decoding performance for each participant (colored dots) and group mean (bars): (c) decoding accuracy (circular correlation between decoded and true location), (d) decoding precision (mean absolute error), and (e) decoding variability (standard deviation of signed decoding errors). Error bars indicate ± sem. (f) Circular correlation between the behavioral memory errors and neural decoding errors across participants. Colored dots represent individual participants; gray bars represent group means; error bars indicate ± sem. ** p < 0.01, permutation test. (g-h) Memory errors plotted against decoding errors, binned into four quantiles (colors, 5 participants per bin), based on decoding error. Gray line represents the best linear fit. Pearson’s r. * p<0.05, permutation test. Other prefrontal regions are plotted in SupFigs 4 & 5.

While we acquired our BOLD data at the isotropic voxel resolution of 0.9mm, we next simulated how our decoding results might have changed with larger more conventional voxel resolutions. To do so, we resampled our data from 0.9mm to 2mm and 3mm isotropic voxel resolutions and recomputed our entire decoding analytic pipeline (see *Methods*). In sPCS, decoding precision decreased linearly with increasing voxel size, but in all other PFC regions, including intFS, there were no systematic effects of voxel size (Sup Fig 7). The effect of decreasing resolution on decoding in areas where decoding was poor to begin with at the highest resolution was not surprising. However, we did not expect decoding to improve at higher resolutions in sPCS because of its map-like retinotopic organization. Even at the resampled 3mm resolution, decoding precision was still better than previous reports from 3T fMRI at 2.5mm isotropic voxels ^31,32,38^ suggesting that SNR improvements due to higher field 7T might improve decoding independent of resolution. While informative as to the impact of resolution, we emphasize that this exercise in resampling underestimates the complexity in the relationships between voxel size, vascular architecture, hemodynamics, and measurement noise ^42–44^.

A key to evaluating the importance of the information encoded in PFC regions is whether or not this information relates to memory behavior. To test for such a relationship, we correlated neural and behavioral memory errors on a trial-by-trial basis. In sPCS, but not intFS, decoding errors during the delay period predicted later errors in memory-guided saccades across participants (p < 0.001; Fig 4f). To further validate this neural-behavioral association estimated from noisy trialwise data, we binned trials by neural decoding error and examined whether decoding errors tracked behavioral errors across bins. This analysis confirmed a robust relationship in sPCS (r = 0.50, p = 0.02; Fig 4g), but no such relationship in intFS (r = 0.07, p = 0.73; Fig 4h). Together, these findings indicate that the readout of delay period activity in sPCS, but not intFS, drives memory behavior.

### Visualization of working memory dynamics and subspace representations

Finally, we examined the temporal dynamics and structural organization of PFC population activity that supports working memory representations. Using voxel receptive field parameters from the pRF models, we projected population activity from cortical space into visual space coordinates to reconstruct activation maps over time (see *Methods*; and ^45^). In sPCS, we observed a bump of activity that emerged at the target’s location beginning 4.5s after the onset of the memory target and remained stable at the remembered location throughout the delay (Fig 5a). To probe the representational geometry of these responses, we conducted principal component analysis (PCA) on time-averaged delay-period BOLD responses to targets segregated into eight position bins (see *Methods*). In the sPCS, the first two PCs accounted for almost 50% of the variance in these BOLD responses, compared to approximately 30% in data that was shuffled (Sup Fig 6). Thus, we projected population activity at each time point into a low-dimensional subspace using the first two PCs (Fig 5b). In sPCS, shortly after the onset of the target, note how the projections spread out over the course of the memory delay (i.e., accounted for increased variance). The topology of the targets was valid and preserved across time. Specifically, the relationships among target locations adhered to their true visual spatial locations suggesting that these two PCS appeared to map onto horizontal and vertical visual space. In contrast, intFS failed to show any clear or structured population activity across the trial (Fig 5c). While there were some hints of information about the target locations in the intFS low-dimensional subspace (Fig 5d), the first two PCs barely accounted for more variance than with the shuffled random data (Sup Fig 7). Moreover, the subspace had several topological errors (i.e., yellow, orange, and red trajectories) and was not stable over the delay period.

**Fig. 5.**
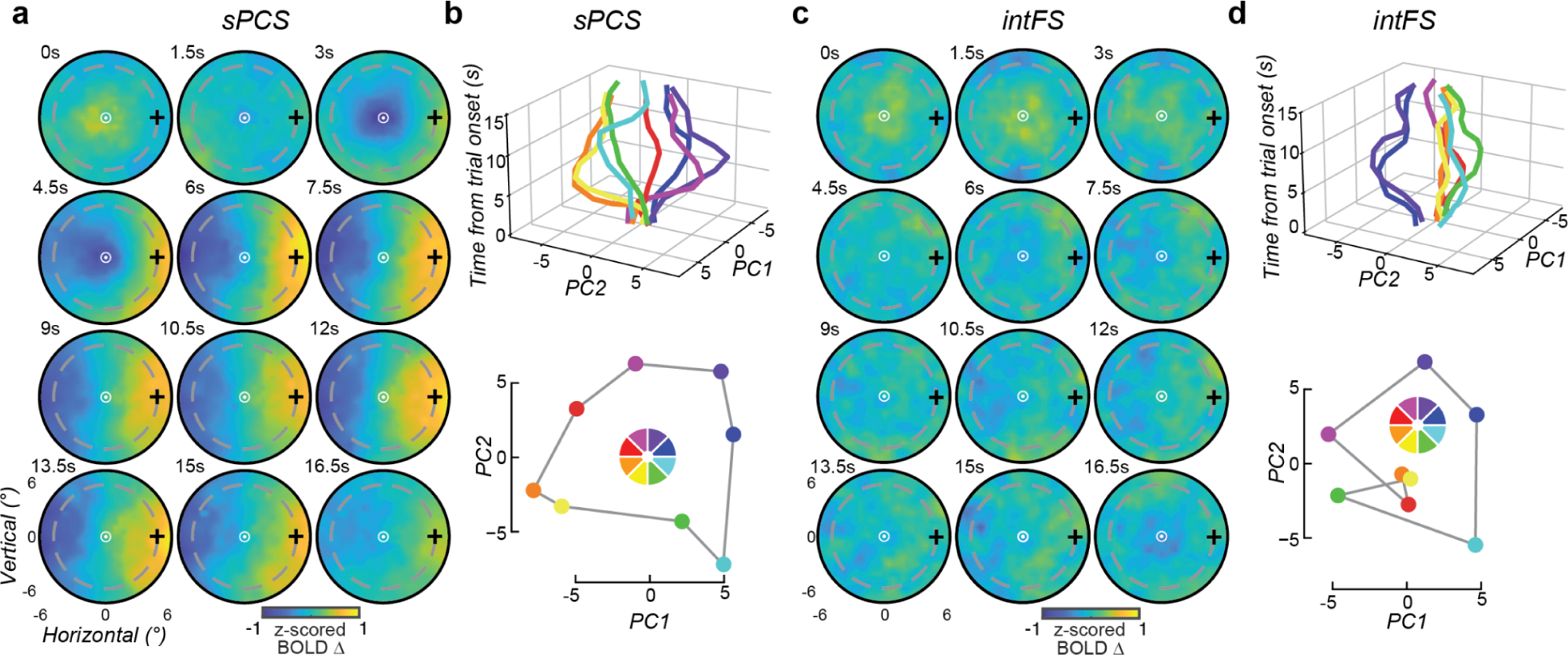
Visualization of working memory dynamics and representational subspaces. **(a)** Reconstructed visual field response maps from sPCS across time. Each map depicts the spatial distribution of voxel activity projected into visual field coordinates, where all trials were rotated to align targets to the 0° (plus sign). **(b)** Top: sPCS responses projected into a low-dimensional subspace defined by the first two principal components (PC1, PC2); the z-axis represents time from trial onset. Each trajectory corresponds to a different target location (color-coded as in the visual field legend). Bottom: Snapshot at 7.5 s after trial onset shows an organized representation of memory targets within the subspace that was topologically valid. **(c, d)** Same as (a, b), shown for intFS. Note the lack of target encoding at the population level and a subspace with several topological errors. Color wheel keys the polar angle of visual space.

## Discussion

We tested the hypothesis that poor spatial resolution of fMRI measurements prevented previous studies from finding working memory related signals in the part of the human PFC homologous to the macaque principal sulcus. Indeed, the clustering of similarly tuned neurons in macaque PFC suggests that a voxel resolution of just above 1mm might be necessary to detect feature selective working memory signals. To address this concern, we scanned the human PFC at 900𝜇m voxel resolution using high-field 7T fMRI to search for behaviorally meaningful, hallmark signatures of working memory storage. In summary, we found that sPCS, and not more anterolateral parcels of the PFC, possessed many of these hallmarks of working memory. Human sPCS 1) contains spatially tuned voxels that form a map of contralateral visual space; 2) activity persists during the memory delay, is contralateralized with respect to the memory target, and is especially prominent in voxels that have strong spatial selectivity; 3) activity patterns can be used to decode fine grain features in working memory; 4) decoding errors predict memory errors on a trialwise basis; and 5) population activity constructs a stable subspace that is topologically yoked to memorized visual space. Below, we discuss how these results impact the functional homologies between the human and non-human primate PFC and theories of working memory.

### High-resolution imaging does not unveil hidden working memory activity in midlateral prefrontal cortex

Our results provide evidence that simply imaging the PFC at high spatial resolution does not reveal previously hidden working memory signals from the intFS–human homolog of macaque principal sulcus–or other parts of the midlateral PFC. In fact, we found no reliable evidence of working memory signals from the intFS. On the one hand, this is surprising as it represents a failure to translate decades of pioneering research into the role of the macaque principal sulcus in working memory. Lesions specifically to the principal sulcus and nearby surrounding convexity cause impairments in spatial working memory ^46–48^, and specifically on the memory-guided saccade task used here ^6,49,50^. Neuronal activity in the principal sulcus and nearby surrounding convexity persist when a memory target is in its receptive field ^1,2^, and specifically during the delay period of memory-guided saccade tasks ^4,25,26,51^. These classic findings deeply shaped neural theories of working memory whereby the PFC is thought to maintain stimulus-specific representations through stable, sustained firing during delay periods ^7–9^.

The animal evidence and coupled theory remain highly compelling, which explains why the lack of similar evidence in human PFC is both surprising and controversial. The first human brain imaging study of spatial working memory, using PET, did not detect increased blood flow in the mid-dorsolateral PFC ^11^. This finding was not unique— subsequent fMRI studies also failed to observe delay-period activity related to spatial working memory in the mid-dorsolateral PFC ^41,52–54^, making such negative results the rule rather than the exception. A later fMRI study from Goldman-Rakic’s lab reported dorsolateral PFC activation, but only when the memory load was increased to five items ^55^. Although the authors argued that fMRI lacked the sensitivity to detect sustained activity associated when maintaining just a single item, it was more likely that when one’s memory capacity was surpassed, control mechanisms housed in PFC were recruited for memory support ^56,57^.

Using fMRI to measure neural activity during spatial working memory tasks—including memory-guided saccades tasks—we consistently find that fMRI is sensitive to single-item working memory representations ^30,34,36,37,58–63^. However, none of these studies reported activity in the human mid-dorsolateral PFC during simple spatial working memory delays. Instead, sustained activity consistently localized to the sPCS and posterior parietal cortex (reviewed in ^16^. Recent studies have shown that powerful machine learning and encoding-decoding techniques can successfully predict the contents of working memory from the fMRI patterns of activity in visual, parietal, and even sPCS in PFC ^30,31,34,61,64,65^. However, in the more anterolateral parts of human PFC no item level information about what is stored in working memory has been reported, except the rare report of orientation information in just a few participants’ right midlateral PFC hemispheres ^19^–a finding which has not been replicated, including in the current study.

Scanning the lateral PFC at submillimeter voxel resolution using high-field 7T did not reveal any hidden working memory information that previous lower resolution 3T studies may have missed. One could always argue that 900𝜇m is still too large and perhaps macrocolumn (∼600𝜇m), microcolumn (∼100𝜇m), or even single PFC pyramidal neuron (∼20𝜇m) resolution is necessary to detect working memory signals. However, this seems unlikely for two reasons. First and despite the non-topographic organization of the mid-lateral PFC, the spatial scale with which similarly tuned neurons are clustered together in and around the principal sulcus suggest that fMRI sampling at a resolution of 1.0 to 1.5mm should be sufficient ^22,23,66^. Second, even if finer imaging resolution were needed, surgical resections of the human lateral PFC only cause deficits in the accuracy of memory-guided saccades when the lesion encroaches into the precentral sulcus ^67–69^. Similarly, temporary disruptions to the human sPCS, but not intFS, with transcranial magnetic stimulation applied during the memory delay impacts the accuracy of memory-guided saccades ^70^. Therefore, not only do discrepancies exist between human and non-human primate findings in terms of delay period neural activity, but also in terms of the impact of lesions to the mid-lateral PFC on memory.

### Hallmark signatures of working memory in the human precentral sulcus

Despite the lack of homology of findings in the more anterolateral PFC, our findings from the superior precentral sulcus (sPCS) represent a remarkable array of hallmark working memory signatures that have largely been restricted to macaque PFC neurons. Note that sPCS likely contains the human homolog of the macaque frontal eye field (FEF). We choose the anatomical term sPCS rather than the function term FEF because in humans we cannot fulfill the electrical stimulation criteria agreed upon by the field to identify FEF neurons in the macaque. Nevertheless, the coarse spatial topography found in the sPCS might correspond to the coarse map of saccade trajectories in FEF ^71,72^.

Delay period activity in sPCS may be the most consistently reported and robust effect across all neuroimaging studies of spatial working memory (e.g., ^11^; reviewed in ^16^). Here at high-field and high resolution, we found that persistent activity was stronger and more lateralized than previous reports at 3T ^30,34,36,37^. Moreover, we discovered that the amplitudes of delay period activity in sPCS were greater in voxels whose spatial tuning were stronger, thus providing an important link between the visual feature selectivity and memory-related persistent activity. Neurons in macaque FEF also show strong and spatially selective persistent activity during the delay of memory-guided saccade tasks ^73–76^.

In fact, when one compares FEF to pre-arcuate to principal sulcus in terms of the strength of delay period activity, spatial selectivity, and prevalence of delay active neurons, neurons in more posterior areas appear to show stronger working memory signals ^4,51,71,77^. In fact, some evidence suggests that a gradient of working memory selectivity exists across these areas in the macaque lateral PFC ^78,79^. Our results from the human PFC do not support such a gradient and suggest that memoranda-specific memory storage is limited to the sPCS. Such a selectivity aligns well with results from both human neuroimaging and lesion studies of the lateral PFC in working memory.

At the population level in sPCS, the patterns of activity contain fine-scale and robust item-level information about the contents stored in working memory. While previous studies at 3T have reported similar results, fine-scale spatial information was absent, effect sizes were small and inconsistent across participants and studies resulting in uncertainty about their meaning ^30–32,38,61,80^. Remarkably, on trials in which the decoded memory target was clockwise, for example, from the actual target, memory-guided saccades generated after the delay tended to also have a clockwise error. Such trialwise coupling between decoded and memory error has only been reported between neuronal spiking in macaque PFC and memory-guided saccade errors ^5^, but are predicted by theory and attractor models of working memory ^8,9^. At high-field and high resolution, we were able to use dimensionality reduction techniques to show that population activity in sPCS formed a low-dimensional subspace of the array of target locations. This subspace was stable throughout the memory delay. Its first two components accounted for half of the variance in the data and was topologically organized such that these two components naturally captured the horizontal and vertical geometry of visual space. Moreover, these results decidedly mirror those found in dimensionality reduction studies of macaque PFC neuronal activity during working memory ^81,82^. Overall, based on the strength of decoding, behavioral coupling, and subspace structure, these results establish the importance of these neural patterns in sPCS and their explicit ties to neural theories that were instead developed to explain how more anterolateral parts of PFC may support working memory. While our results in human PFC are rare, similar findings have been reported from studies of the parietal and visual cortices ^15,30,31,83,84^. Given that working memory signals are so broadly distributed, future research should focus on how these areas both differ and coordinate to support working memory.

To summarize, decoding errors in sPCS predicted behavioral performance on a trial-by-trial basis, confirming the behavioral relevance of these representations. Furthermore, dimensionality reduction revealed that the population response in sPCS occupied a low-dimensional subspace that preserved the topological structure of visual space—an organizational feature suggestive of systematic, stable neural encoding. These findings align with a growing body of evidence that implicates posterior regions of the lateral prefrontal cortex, particularly the PCS and surrounding sulci, in spatial working memory storage.

### Functional considerations beyond working memory storage

While canonical theories of the PFC’s role in working memory have largely focused on memory storage ^7,10^, our results have broader implications for regional and functional specialization. Without using the memory-guided saccade task to assess working memory we could not make comparisons with the results from macaque electrophysiology studies using the same task. However, it is possible that the delay period activity we measured in sPCS could be partially due to sustained spatial attention or saccade planning functions, as well as memory storage ^34^. On the one hand, this limits our ability to precisely specify the process involved. On the other hand, and opposed to short term memory, working memory is the process by which mnemonic information guides our natural behaviors ^85,86^, including actions like gaze shifts. Moreover, population activity in human sPCS ^34,87,88^, and macaque FEF ^89,90^, may encode a prioritized map of visual space whose read-out can support several critical cognitive functions beyond just working memory ^91^. For instance, while the sPCS/FEF may not contain the specific motor metrics used to generate saccades, readout of the eye-centered spatial coordinates stored in sPCS/FEF by subcortical structures like the superior colliculus and brainstem saccade generator could be used to plan saccades ^74,92^. Similarly, readout by visual areas in extrastriate cortex could be the mechanism by which neurons whose receptive fields match the prioritized portion of visual space are selectively boosted, causing the effects of spatial attention ^93,94^. Thus, the same priority maps in sPCS could efficiently support working memory, spatial attention, and oculomotor planning.

### Potential reasons for lack of homology between species

In light of these new findings, let us reason about what might explain why these differences between humans and non-human primates exist and what are the implications for working memory theory. Perhaps the PFC architectures in the two species, which diverged 25-30 millions years ago, evolved differently to support spatial working memory. Although this would require that we amend our neural theories of working memory, they would not require a major overhaul as the precise mechanisms (e..g, persistent activity; feature-distributed population coding; recurrent neural networks, etc) would need no change. Moreover, it is highly unrealistic to think that no differences would exist between the two species. Anatomically, the macaque principle sulcus evolved into three tertiary sulci, the intermediate frontal sulci (intFS), that extend along the anterior-posterior axis of the middle frontal gyrus ^13,14^. Functionally, in the precentral sulcus there are at least three visual field maps in humans ^67^, and likely only one in macaque arcuate sulcus corresponding to the FEF ^71,95^. Similarly, there are at least six visual field maps in human parietal cortex ^67,96,97^ compared to two in the macaque parietal cortex ^98,99^. Nonetheless, the canonical theory of working memory may only require a small change to accommodate these new findings– a small posterior shift of the locus of working memory.

While the type of neuronal signals electrophysiology and BOLD fMRI are sensitive to differ, there are reasons why these differences are unlikely to be the culprit here. There are no reasons to think that BOLD is less sensitive to measurements from the precentral sulcus compared to the shallower intermediate frontal sulci that are closer to the MRI’s RF coils. Plus, BOLD and spiking should be highly correlated during memory delays when neuronal activity persists; the correlation between BOLD and spiking only reduces when neuronal spiking adapts ^20,100^. Moreover, perhaps differences in the subjective difficulty of the single item task might play a role ^38^. Memorizing a single target’s location might be considered easier to humans compared to macaques, but offsetting this concern, macaques have orders of magnitude more practice ^101,102^. In humans, however, training itself has little if any impact on item specific representations in lateral PFC beyond a single training session ^103^. What about motivational factors that clearly differ? In macaque studies, rewards are needed to enforce behavior, while humans seem intrinsically motivated to minimize errors on working memory tasks ^38,104^. Finally, if increased memory loads are required to activate anterolateral PFC ^55,105^, could this suggest that control rather than storage processes are the primary function of the PFC ^32,57,87,106^. Executive control demands are rather weak in the memory-guided saccade task, which might explain why we observed little delay period activity in the more anterior portions of the PFC. While none of these measurement, task, process, or motivational factors currently figure into our neural theories of working memory, they do not seem likely to explain key failures of translating the macaque PFC results in humans.

## Conclusions

Our findings challenge the canonical view that the midlateral PFC serves as the primary locus of working memory storage in both humans and macaques. In humans, we show that persistent, stimulus-specific representations supporting working memory are localized to sPCS, rather than the more anterior inFS. Population activity in the sPCS might best be conceptualized as a prioritized map of space used for a variety of spatial cognitive tasks (e.g., working memory, attention, and intention). This cross-species dissociation highlights the need for caution when translating neural mechanisms from non-human primate models to the human brain, and calls for a careful reevaluation of long-standing animal models of working memory that may have overemphasized the centrality of the midlateral PFC. At the same time, our results affirm that the core computational mechanism of working memory—stimulus-specific persistent activity within spatially organized neural populations—is conserved across species. That this mechanism is implemented in different anatomical substrates suggests evolutionary adaptations that may support the expanded behavioral and cognitive repertoire of humans, particularly in domains such as abstract reasoning, planning, and flexible control. Understanding the extent and nature of these cross-species differences will be critical for building unified, biologically grounded models of working memory and for assessing the translational nature of animal models to human cognition.

## Acknowledgements

This work was supported by National Institutes of Health Grants R01 EY033925 to CEC and R01 EY034118 to KNK and CEC. We thank the University of Minnesota’s Center for Magnetic Resonance Research for support.

## Methods

### Participant details

The study was advertised to the University of Minnesota community. Potential participants were contacted for a Zoom interview to explain the nature of the study and to screen them for eligibility. Based on the Zoom interviews, we invited 10 individuals to participate in an initial 7T fMRI screening session. Of these, five were invited to participate in the full experiments (one of these was an author). All participants were right-handed individuals with no known cognitive deficits nor color blindness and had normal or corrected-to-normal vision. The experiments were conducted with informed written consent of all participants, and the experimental protocol was approved by the University of Minnesota Institutional Review Board. Participants were compensated at a rate of $30 per hour, plus performance bonuses. Our experimental approach emphasizes collecting extensive data from a small set of participants whom we were confident would each provide high-quality data, rather than collecting a small amount of data from a larger, less selective sample.

### Memory-guided saccade (MGS) task

Each trial began with the brightening of a central fixation point, serving as a cue to alert the participant to the impending appearance of a target. One second later, a working memory (WM) target (a light gray dot, 0.325° diameter) was presented for 500 ms, followed by a delay period of 12,000 ms. Participants were instructed to remember the location of the target while holding their gaze at the fixation point at the screen center throughout the delay. After the delay period, the fixation point disappeared, serving as a response cue, instructing participants to make a saccadic eye movement to the remembered location. 700 ms after the onset of the response cue, a feedback stimulus (a white dot) was presented at the target location for 800 ms. Participants were instructed to make a corrective saccade to the feedback dot before returning their gaze to the screen center. The intertrial interval was pseudo-randomly chosen to be 6, 9, or 12 s, during which the fixation point dimmed. Each participant completed 320 or 352 trials across two 1.5-hr scanning sessions on separate days. Each session included 10 or 12 runs, each run consisting of 16 trials. Target locations were evenly distributed across circular space in 8 locations positioned at an eccentricity of 5.5°. A full set of 8 locations were shown in random order, followed by another full set of 8. Each trial included a spatial jitter, ranging from –5.625° to 5.625° (uniform random sampling). Participants were allowed to take a break between runs.

### RSVP retinotopic mapping task

Each participant was scanned for one 1.5-hour retinotopic mapping session to identify regions-of-interest (ROIs) and to estimate each voxel’s population receptive field (pRF), following established procedures in Mackey et al. ^33^. During each retinotopy run, participants performed an attention-demanding object image rapid serial visual presentation (RSVP) task. In the RSVP task, while maintaining fixation on a central cross, participants covertly tracked a moving stimulus bar composed of six equally sized color images of household objects. The bar swept across 5×9° of the visual field in twelve discrete 2.6-s steps. Images were presented for a certain duration (ranging between 150–600 ms) and then replaced with new ones. In each sweep, participants reported whether the target object image was present among the six images via button press. The participants received online feedback: if they correctly responded within 800 ms of the presentation of the target object, the fixation point briefly turned green. The target image was pseudo-randomly chosen for each run and was shown at the start of each run to help participants get familiar with it. The presentation duration of the images comprising the bar was adjusted based on participants’ performance via a staircase procedure; the bar’s sweep speed remained fixed. Participants were required to fixate on the center of the screen throughout the task. Their fixation was monitored online and confirmed via eye tracking. Participants performed 8 bar sweeps per run.

### Stimuli display and eye tracking setup

We presented stimuli on a Cambridge Research Systems BOLDscreen 32 LCD monitor (1920 × 1080 pixels at 120 Hz), positioned at the head of the 7T scanner bed and flush against the scanner bore. Participants viewed the monitor via a mirror mounted on the RF coil. The total viewing distance from the eyes to the monitor (via the mirror) was approximately 184 cm (5 cm from eyes to mirror + 178.5 cm from mirror to monitor). The size of the full monitor image was 69.84 cm (width) × 39.29 cm (height). At the beginning of each scan session, we positioned the monitor, the participant’s head, and RF coil in the same location relative to the scanner.

A Mac computer controlled stimulus presentation using code written in PsychoPy for the pRF experiment and in Psychophysics Toolbox 3.0.14 for the MGS experiment. We recorded behavioral responses using a button box (Current Designs). In all scan sessions, we performed eye-tracking using an EyeLink 1000 system (SR Research) with a custom infrared illuminator mounted on the RF coil. We tracked the left eye at 500 Hz using the Pupil-CR centroid mode. At the beginning of each session and as necessary between runs, we calibrated the eye tracker using a 9-point routine for the MGS experiment and a 5-point routine for the pRF experiment. We monitored gaze data and adjusted pupil/corneal reflection detection parameters as needed during and/or between runs. We also collected physiological data using a pulse oximeter and a respiratory belt (stock Siemens equipment). We taped the oximeter to the participant’s left index finger and secured the respiratory belt snugly around the torso. We do not analyze the physiological data in this study.

### MRI data acquisition

MRI data were collected at the Center for Magnetic Resonance Research at the University of Minnesota. Anatomical (T1- and T2-weighted) data were collected with a 3T Siemens Prisma scanner and a standard Siemens 32-channel RF head coil. Functional data (T2*-weighted) were collected using a 7T Siemens Magnetom actively-shielded scanner and a single-channel-transmit, 32-channel-receive RF head coil (Nova Medical, Wilmington, MA). To mitigate head motion, we used standard MR-compatible foam padding on the back of the head, along with additional neck and ear padding.

We collected several anatomical data at 3T (T1 and T2). The motivation for collecting data at 3T was to ensure acquisition of T1 volumes with good gray/white-matter contrast and homogeneity, which is difficult to achieve at ultra-high fields. To increase contrast-to-noise ratio and enable the ability to assess reliability, we acquired several repetitions of T1-weighted and T2-weighted volumes. For each participant, we collected 3 scans of a whole-brain T1-weighted MPRAGE sequence (0.8-mm isotropic resolution, TR = 2,400 ms, TE = 2.22 ms, TI = 1,000 ms, flip angle 8°, bandwidth 220 Hz per pixel, no partial Fourier, in-plane acceleration factor (iPAT) 2, TA = 6.6 min per scan) and 2 scans of a whole-brain T2-weighted SPACE sequence (0.8-mm isotropic resolution, TR = 3,200 ms, TE = 563 ms, bandwidth 744 Hz per pixel, no partial Fourier, iPAT 2, TA = 6.0 min per scan).

We collected functional data and associated fieldmaps at 7T, including 1 session for the pRF experiment and 2 sessions of the MGS experiment. In the pRF session, functional data were collected using gradient-echo EPI at 1.8 mm isotropic resolution with whole-brain (including cerebellum) coverage (84 axial slices, slice thickness 1.8 mm, slice gap 0 mm, field-of-view 216 mm (FE) × 216 mm (PE), phase encode direction anterior-to-posterior, matrix size 120 × 120, TR = 1,600 ms, TE = 22.0 ms, flip angle 62°, echo spacing 0.66 ms, bandwidth 1,736 Hz per pixel, partial Fourier 7/8, iPAT 2, multi-band slice acceleration factor 3). In the MGS session, functional data were collected using gradient-echo EPI at 0.9 mm isotropic resolution with partial brain coverage (60 slices, slice thickness 0.9 mm, slice gap 0 mm, field-of-view 135 mm (FE) x 133.2 mm (PE), phase-encode direction right-to-left, matrix size 150 x 148, TR 2100 ms, TE 27.4 ms, echo spacing 1.05 ms, bandwidth 1076 Hz/pixel, partial Fourier 6/8, in-plane acceleration factor (iPAT) 2, multiband slice acceleration factor 2). The flip angle for the functional data was set based on empirical flip angle measurements, and ranged around 42-55 degrees across participants. In addition to the EPI scans, we also collected dual-echo fieldmaps for post-hoc correction of EPI spatial distortion (same overall slice slab as the EPI data, pRF fieldmaps: 2.2 mm x 2.2 mm x 3.6 mm resolution, 42 slices, TR 510 ms, TE1 8.16 ms, TE2 9.18 ms, flip angle 40°, bandwidth 350 Hz/pixel, partial Fourier 6/8, TA 1.3 min/scan, MGS fieldmaps: 2.2 mm x 2.2 mm x 2.7 mm resolution, 20 slices). Fieldmaps were periodically acquired over the course of each scan session to track changes in the magnetic field.

Note that the 0.9-mm protocol has a limited field of view. We used care in placing the slice slab to ensure coverage of the frontal brain regions. We show a sample slice screenshot to give a sense of where this was (see Supplementary Fig 10). Note that the field-of-view was relatively tight in the left-right dimension (hence, a small amount of wraparound artifacts exist in the most inferior slices). We optimized the parameters of the pulse sequence to maximize signal-to-noise ratio while delivering high spatial resolution measurements.

### fMRI data preprocessing

The fMRI data were preprocessed using a custom pipeline (based on procedures from ^107^). Pre-processing results were carefully visually inspected to ensure quality control.

T1- and T2-weighted volumes were corrected for gradient nonlinearities using a custom Python script (https://github.com/Washington-University/gradunwarp) and the proprietary Siemens gradient coefficient file retrieved from the scanner. The T1 and T2 volumes acquired for a given participant were then co-registered (within modality). This was accomplished by co-registering all T1 (and T2) volumes (rigid-body transformation; correlation metric). In the estimation of registration parameters, a manually defined 3D ellipse was used to focus the cost metric on brain tissue. Multiple acquired volumes were cubic interpolated to implement the co-registration, and then averaged (within modality) to increase contrast-to-noise ratio.

The T1-weighted volume was processed using FreeSurfer version 7.4.1 with the-hires option to take advantage of the 0.8-mm resolution. We also provided the co-registered and averaged T2 to FreeSurfer and used the -T2pial option. We used an expert file to specify a larger number of inflation iterations (50). As part of FreeSurfer processing, the T2-weighted volume was aligned and resampled to match the T1-weighted volume. Using mris_expand, we generated six cortical surfaces positioned equally spaced between 10% and 90% of the distance between the pial surface and the boundary between gray and white matter.

In the first stage of fMRI preprocessing, the fMRI data were pre-processed by performing one temporal resampling (cubic interpolation) to correct for slice time differences. A cubic interpolation of each voxel’s time-series data in each run was performed. This interpolation corrected differences in slice acquisition times and also upsampled the data (in the same step) to 1.3 s for the pRF experiment and 1.5 s for the MGS experiment. Fieldmaps were prepared as described in ^108^ and were linearly interpolated over time, producing an estimate of the field for each fMRI volume acquired. Temporally resampled fMRI volumes were undistorted based on the field estimates. The undistorted volumes were used to estimate rigid-body motion parameters using the SPM5 utility spm_realign. One spatial resampling (cubic interpolation) of the temporally resampled volumes was used to correct for the combined effects of head motion and EPI distortion.

Then, we aligned the mean pre-processed fMRI volume to the T2-weighted volume (prepared by FreeSurfer) using a small amount of nonlinear warp (SyN) as implemented by ANTs 2.1.0. (Nonlinearity was necessary to compensate for acquisition of the anatomical data and functional data on different scanners.) A separate registration was performed for the mean volume obtained for each fMRI run. The parameters of ANTS were tuned to maximize registration quality.

In the second stage, we coupled the estimated nonlinear warp with the first-stage transformations to produce a final preprocessed fMRI volume in register with the anatomy. Two versions of the functional data were prepared. For the pRF data, we used a 1.8-mm grid parallel to the FreeSurfer anatomical volume (upsampled temporal-resolution, 1.3 s). For the MGS data, we used a 0.9-mm grid within a box conformed to cover the extent of the partial-brain coverage data (upsampled temporal-resolution, 1.5 s). The final result of fMRI pre-processing was volumetric fMRI time-series data in participant-native space (registered to the anatomy).

### Identification of regions of interest (ROIs)

We first averaged each voxel’s time series across all retinotopy runs and fit a population receptive field model with compressive spatial summation to the resulting BOLD time series for each participant ^109,110^. We then visualized the pRF model results on the cortical surface by projecting best-fit polar angle and eccentricity parameters with variance explained ≥ 10% onto each participant’s inflated surfaces using Connectome Workbench. Next, we manually defined regions of interest based on visual inspection, following established criteria such as polar angle reversals and foveal representations ^33^. Finally, we projected the ROIs back into volume space to select voxels for analysis. In the main text, we focus on data from the superior precentral sulcus (sPCS), but see Supplementary Information for analyses involving the inferior precentral sulcus (iPCS). Moreover, despite that sPCS is actually composed of two maps, here we simply treat them as a single map as in previous studies ^30–32^.

In addition to the functionally defined sPCS based on pRF parameters, we defined anatomical ROIs in the superior frontal gyrus (sFG), superior frontal sulcus (sFS), middle frontal gyrus (mFG), inferior frontal sulcus (iFS) and intermediate frontal sulcus (intFS) using each participant’s FreeSurfer parcellation. For each anatomically defined ROI, we excluded voxels overlapping with any functionally defined ROI. Here, we focus on data from the intFS because those set of tertiary sulci are thought to be the human homolog of the macaque principle sulcus based on cross species concordances in cytoarchitecture and resting state fMRI data ^13^). See Supplementary Information for results from the other anatomical regions.

### GLM analysis of MGS task

We performed a general linear model (GLM) analysis of the pre-processed time series data from the MGS experiment to estimate the magnitude of trial-wise BOLD responses for subsequent decoding analyses. The GLM was designed to yield a separate beta estimate for each trial’s delay period by including a distinct regressor per trial. We implemented this trial-wise GLM in AFNI using 3dDeconvolve. For four participants, the design matrix included 320 delay-period regressors, and for one participant, it included 352, corresponding to the number of total trials. Each regressor modeled the BOLD response during the 6-second WM delay period (4–10 s after delay onset) using a boxcar function time-locked to delay onset and convolved with a canonical hemodynamic response function (HRF). We implemented the HRF using AFNI’s “GAM(p,q,d)” function with default shape parameters (p = 8.6; q = 0.547) and a duration parameter (d) adjusted to match the 6-second delay epoch.

In addition to the trial-specific delay-period regressors, the GLM included two additional regressors, one coding for stimulus onset and one coding for response period, as well as six motion parameters from the preprocessing step. The model was estimated using ordinary least squares, yielding a unique beta value for each trial’s delay period. These trial-wise beta estimates were extracted from each voxel within each ROI and used to train and test the Bayesian decoder (see below, TAFKAP decoding). We also included a contrast comparing the average delay-period activity to baseline; the resulting voxelwise t-values served as estimates of delay-period activation strength and were used to identify voxels with robust WM-related activity.

### TAFKAP decoding

We decoded the content of WM on a trial-by-trial basis using a Bayesian generative model known as TAFKAP (The Algorithm Formerly Known as Prince, https://github.com/jeheelab/TAFKAP), previously described by ^39^ and ^40^. For each voxel and each trial, we used the trialwise beta values from the GLM, which reflected the estimated delay-period activity, as input to the model. While the ROIs were first defined by different means - sPCS/iPCS were defined by pRF mapping and intFS and other PFC ROIs were defined by anatomy - we then applied a uniform functional criteria to select voxels within the ROIs for decoding and all other analyses. Specifically, we matched the number of voxels by selecting the 1,000 voxels within each ROI that exhibited the strongest delay-period responses, as determined by their t-values from the GLM contrast comparing delay-period activity to baseline. These selected voxels were used both to train the model, and to decode the data from held-out trials in the test set.

For each participant and each ROI, we trained the model using a leave-one-run-out cross-validation procedure. During training, TAFKAP used bootstrap aggregation (bagging) to estimate the free parameters of the generative model: trials in the training set were resampled with replacement multiple times, and a separate set of parameter estimates were obtained from each resampled dataset using ordinary least squares. To decode each trial in the held-out test run, the model applied Bayes’ rule to invert the learned generative model and generate a posterior distribution over stimulus locations. This process was repeated for each set of parameters estimated from the resampled training data. The resulting posterior distributions were averaged to yield a final decoded posterior distribution per trial, from which we computed the circular mean as the decoded stimulus location. Additional details on the model fitting and decoding can be found in ^39,40^ and prior applications of this approach have been reported in ^31^ and ^38^.

To ensure our decoding results are not depending on how we define ROI and how we select voxels, we performed two additional control analyses. First, we applied a different functional criteria for voxel selection in addition to delay-period responses. We selected 1000 voxels in each ROI whose pRF model fit surpassed a common threshold (pRF R² ≥ 10%). We observed results indistinguishable from our main results across ROIs, suggesting that the WM signals we decoded are not driven by our voxel selection strategy (see Supplementary Figure 6). Second, we used a purely anatomically defined sPCS (from the Freesurfer S_precentral_sup_part parcel) and again these results were indistinguishable from the functionally defined sPCS (t=1.76, p=0.16), suggesting that the use of pRF mapping to define ROI cannot explain our results. Together, these additional analyses confirm that our decoding results are robust and invariant to the ROI definition and voxel selection strategy.

To assess decoding performance at more conventional fMRI resolutions, we resampled the pre-processed time-series data from 0.9-mm to 2-mm and 3-mm isotropic voxel sizes using an ideal Fourier filter with cubic interpolation, and repeated the full TAFKAP decoding pipeline at each resolution.

### Behavioral data analysis

For the MGS experiment, we used gaze position as a behavioral measure of WM performance. We preprocessed raw gaze data using fully automated procedures implemented in iEye_ts (https://github.com/clayspacelab/iEye). These procedures removed blinks, corrected for within-run drift, recalibrated gaze data on a trial-by-trial basis, identified memory-guided saccades, and flagged trials for exclusion based on predefined criteria.

We defined blinks as 100 ms before and after any period when pupil size dropped below the 0.1th percentile of the within-run pupil size distribution (run duration: 396 s). We computed gaze velocity from smoothed gaze time courses (Gaussian kernel, 5 ms standard deviation) and defined saccades based on a velocity threshold of 30 deg/s, a minimum duration of 7.5 ms and a minimum amplitude of 0.25°. We defined periods between saccades as fixations. We corrected for drift on each trial based on the fixation position during the fixation and delay period before the response cue. To recalibrate gaze traces on a trial-by-trial basis, we identified the final fixation to the known target location during the 800 ms feedback period, when participants were instructed to fixate the re-presented target, and used this fixation to recalibrate the X and Y traces for that trial within a run.

We quantified WM error based on the endpoint of the initial saccadic eye movement toward the remembered location, referred to as the ‘primary saccade’. Primary saccades were defined as those with an amplitude greater than 2°, a saccade duration less than 850 ms and occurring after the response cue. If participants did not make a corrective saccade before the feedback stimulus appeared, we treated the primary saccade endpoint as the final response. To be included in the analysis, the primary saccade had to both initiate and terminate within the designated response period.

We excluded trials based on two criteria: (1) failure to identify a valid primary saccade, or (2) excessive error for primary saccade, defined as a deviation greater than 10° of visual angle from the target location. For fMRI model estimation (GLM and TAFKAP decoding), we included all trials regardless of behavioral performance to ensure balanced sampling of spatial locations. However, we restricted subsequent analyses, such as quantifying the decoding performance and relating decoded estimates to behaviors, to trials with reliable behavioral performance. To quantify participants’ behavioral memory error, we calculated the signed angular difference (in polar angle) between the reported location, as indicated by the primary saccade endpoint, and the true WM target location.

### Statistical analysis

We applied permutation-based statistical procedures. To generate a null distribution, we randomly permuted the target locations and re-performed the full TAFKAP decoding analysis 500 times. For each participant and each ROI, we compared the distribution of signed decoding error obtained from the real data against the null distribution derived from the permuted data (Figure 4b). We further quantified decoding performance using three metrics (Figure 4c-e): (1) circular correlation between the decoded and true target locations (decoding accuracy), (2) mean absolute decoding error (decoding precision), and (3) standard deviation of decoding error (decoding variability). For each participant and ROI, we compared the observed metric values to the corresponding null distribution obtained from randomly shuffling the target locations and then recalculating the metric for 2000 times. To assess statistical significance at the group level, for each participant, we randomly draw one metric value from their respective null distribution of 2000 values, and calculate the average of these randomly drawn values from all participants. We repeated 10000 times to construct null distributions for the averaged decoding metrics at the group level.

To relate neural decoding performance to behavior, we conducted two complementary statistical analyses. First, we tested trial-by-trial correlations by computing the circular correlation between decoding error and memory error for each participant, then averaging the resulting correlation coefficients across participants (Figure 4f). To evaluate statistical significance, we repeated this analysis using decoding errors from the same permutation procedure (i.e., target locations shuffled), generating a null distribution of average correlation coefficients across 500 iterations. We computed two-tailed p-values as the proportion of permutation yielding correlation coefficients more extreme than the actual correlation coefficient. Second, we performed a binned-correlation analysis by sorting trials into four bins based on increasing decoding error and computing the mean memory error within each bin. We pooled data across participants (four data points per participant), subtracting each participant’s mean to remove between-subject variance. Pearson correlations were computed on the pooled, mean-centered dataset. For visualization purposes (Figure 4g), we re-added the grand means back to each participant’s data. To determine statistical significance, we compared the correlation coefficient to the null distribution obtained by permuting the data labels in the pooled dataset 1,000 times.

### Visualizations of WM representations

We used each voxel’s pRF parameters to visualize how neural populations represented remembered spatial locations. To compute activation maps (Figures 5a & c), we defined a 2D spatial grid centered on fixation, sampling visual space in 0.5° steps across a ±6° range in both horizontal and vertical directions. For each voxel, we modeled its spatial sensitivity as a bivariate circular Gaussian centered at its estimated pRF location (*x_0_*, *y_0_*), with the width determined by its pRF size (σ). The Gaussian weight at each grid point (*x_i_*, *y_i_*) was computed as:

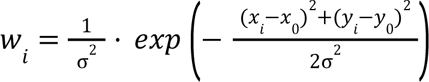

This formulation ensures that voxels with narrower pRFs (i.e., smaller σ) contribute more localized and higher-amplitude spatial weights, while broader pRFs contribute more diffusely and with lower gain. The resulting weight matrix had dimensions of *n_gridpoints_* × *n_voxels_*, reflecting the voxel’s sensitivity to each visual field location. To ensure fair contribution across grid points, weights were normalized at each grid point across voxels: for each location, the weights from all voxels were summed, and each voxel’s weight was divided by this sum. This normalization ensured that the total weight across voxels remained constant at every visual field location. For each grid point, we computed activation as a weighted sum of voxel responses within each ROI:

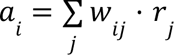

where *r_j_* denotes the response of voxel *j*, and *a_i_* is the reconstructed activation at spatial location *i*. We included only voxels whose pRF model explained at least 10% of the variance during the retinotopic mapping session. To generate an activation map for a given time window in each ROI, we first computed a trial-level activation map for each ROI, then rotated each trial’s map so that the remembered target location aligned to 0° polar angle. These aligned maps were averaged across all trials within each participant, then across participants. Finally, to facilitate visualization of relative spatial patterns, we subtracted the grand mean from the final group-level activation map. Further details on the implementation of circular Gaussian function can be found in a prior application ^45^.

### Representational subspaces

We used principal component analysis (PCA) to define low-dimensional subspaces capturing the representational structure of remembered target locations. For each ROI, we constructed a data matrix X of size of *n_stimuli_* × *n_voxels_*, where each row corresponds to the mean voxel activity pattern across all trials for a given target location. We included the same set of voxels used in the decoding analyses, those exhibiting strong delay-period activity. Prior to applying PCA, we mean-centered each column of X, such that each voxel’s activity was demeaned across stimulus conditions. We concatenated the voxel activity patterns across participants within each ROI and applied PCA on the participant-aggregated matrix. This approach allowed us to capture shared structure in the representational geometry across individuals while preserving inter-subject variability in voxel-level response patterns.

To define the representational subspaces, we first removed temporally varying information by averaging voxel activity across all time points within 3–9 seconds of the delay period. We then performed eigendecomposition on the covariance matrix of the resulting averaged data to extract the principal components (PCs). We defined the subspace using the first two PCs, corresponding to the top two eigenvalues, resulting in a reduced weight matrix W of size *n_voxels_* × 2 (See Supplementary Figure 9 for scree plots and the variance explained by the first two PCs). To visualize the temporal dynamics of population responses within this subspace, we projected the data at each time point during the delay period into the 2D subspace. For each time point, we computed T=XW, where X is the voxelwise activity matrix at that time point. The resulting matrix T, of size: *n_stimuli_* × 2, was the projection of the voxel activity pattern of each stimulus location in the subspace defined by the top two PCs. These projections captured the evolving trajectory of stimulus-location-specific representations in the low-dimensional space over time. Additional details on this eigendecomposition approach can be found in a prior application by ^45^.

## Supplementary Figures

**Supplementary Figure 1.**
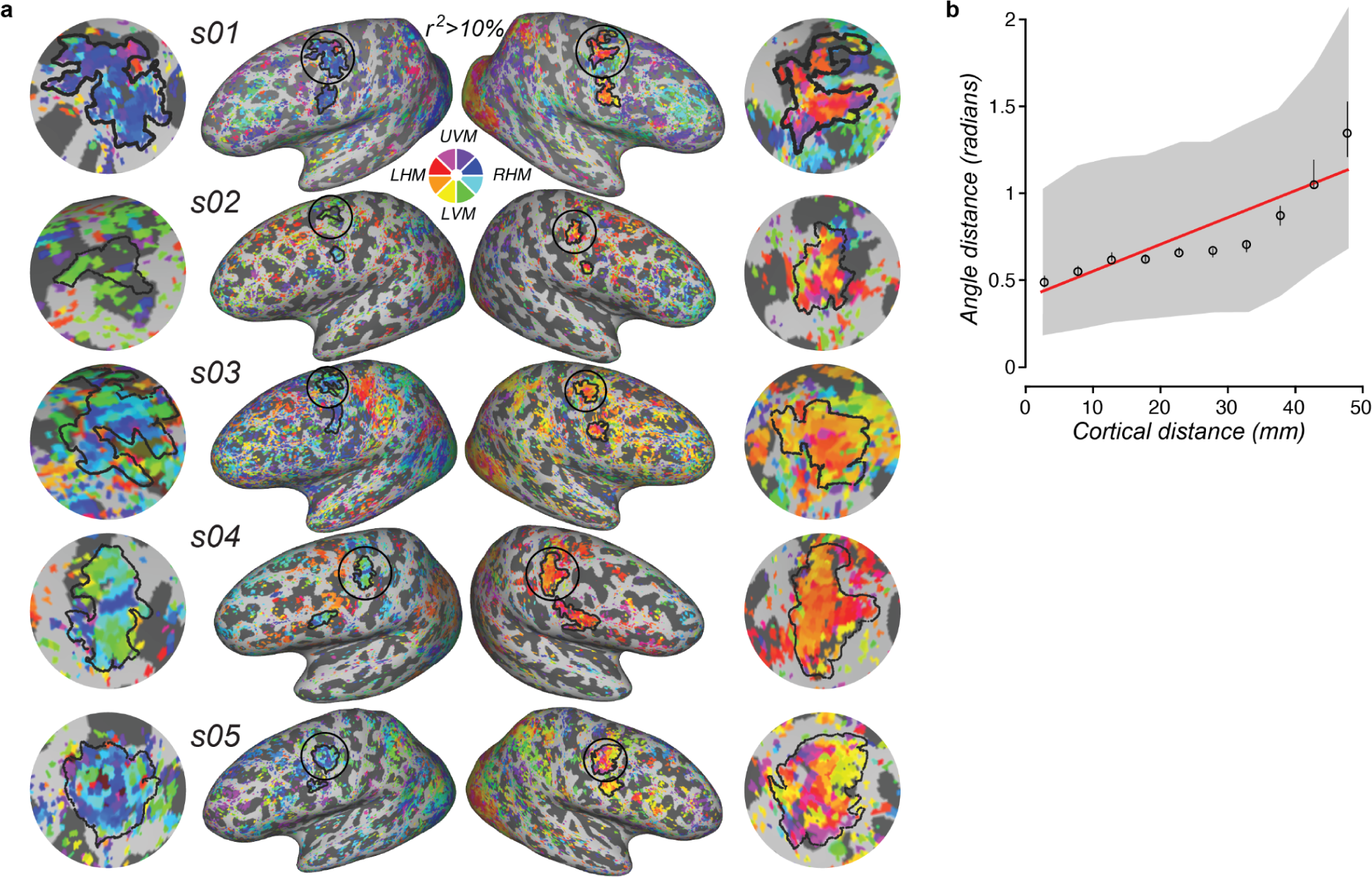
(a) Spatial selectivity and visual field maps in all subjects, related to figure 2. Using population receptive field (pRF) mapping, we modeled the receptive field properties of each voxel to quantify the spatial selectivity of voxels and define retinotopic maps in frontal cortex for each of the participants. Retinotopic maps of contralateral visual space can be seen in superior precentral sulcus (sPCS; putative homolog of macaque frontal eye field; black outlines) for all participants. Close-up views of sPCS are shown on the sides. The color wheel keys the polar angle of the visual field, which is overlaid on the cortical surface thresholded by variance explained (>0.1). (b) Absolute polar angle difference between two voxels plotted against cortical distance resulted in a significant correlation (r = 0.102, p < 0.001). We divided the cortical distance from 0 to 50 mm into ten bins. The circles are the medians of each bin, error bars are the 95% confident interval of this median, gray shade denotes the 25th and 75th percentile of each bin, and the red line is the linear fit across the medians.

**Supplementary Figure 2.**
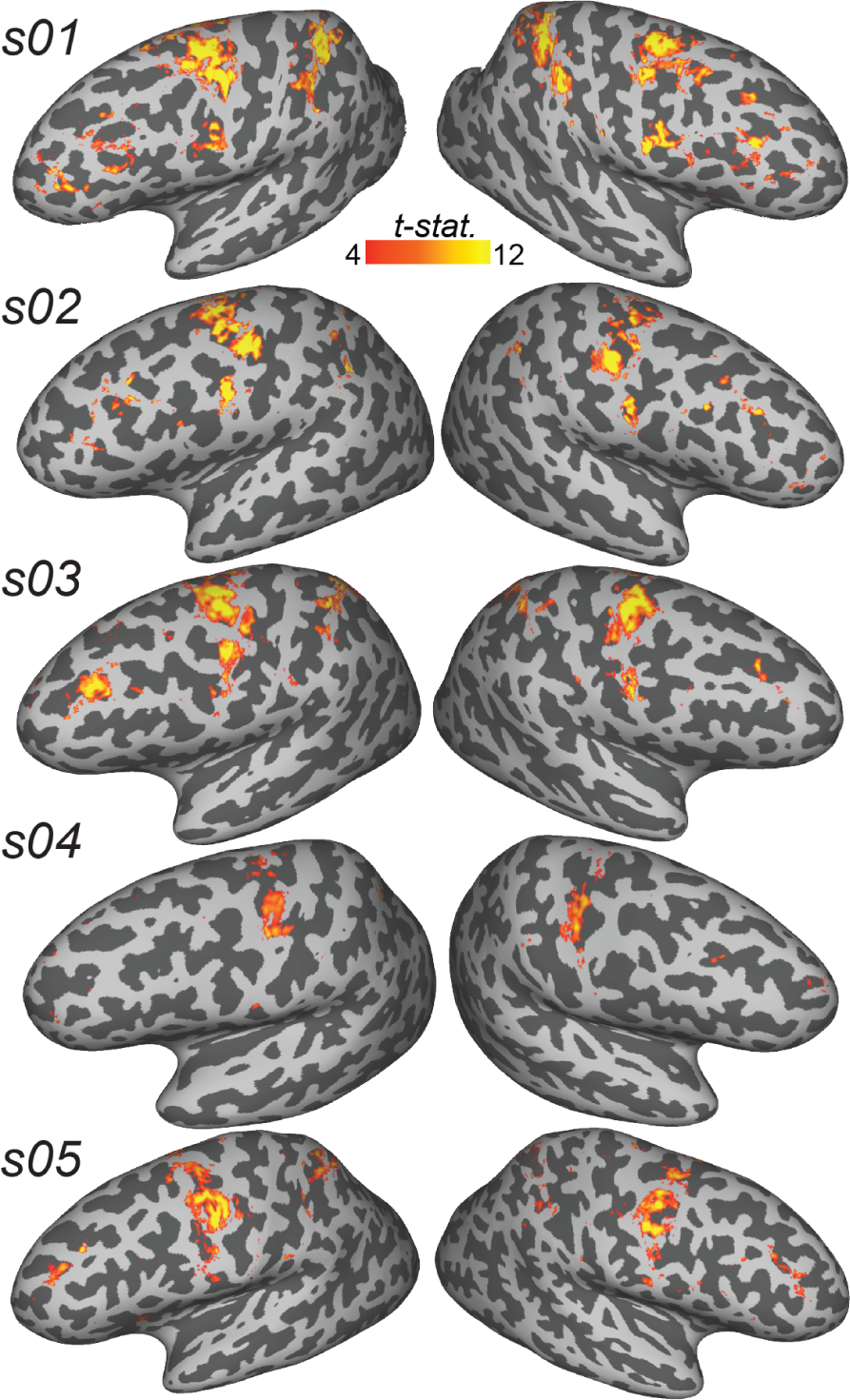
Delay period activity in all participants, related to Figure 3. Statistical maps of delay period activity in each individual participant overlaid on cortical surface, thresholded at t > 4.

**Supplementary Figure 3.**
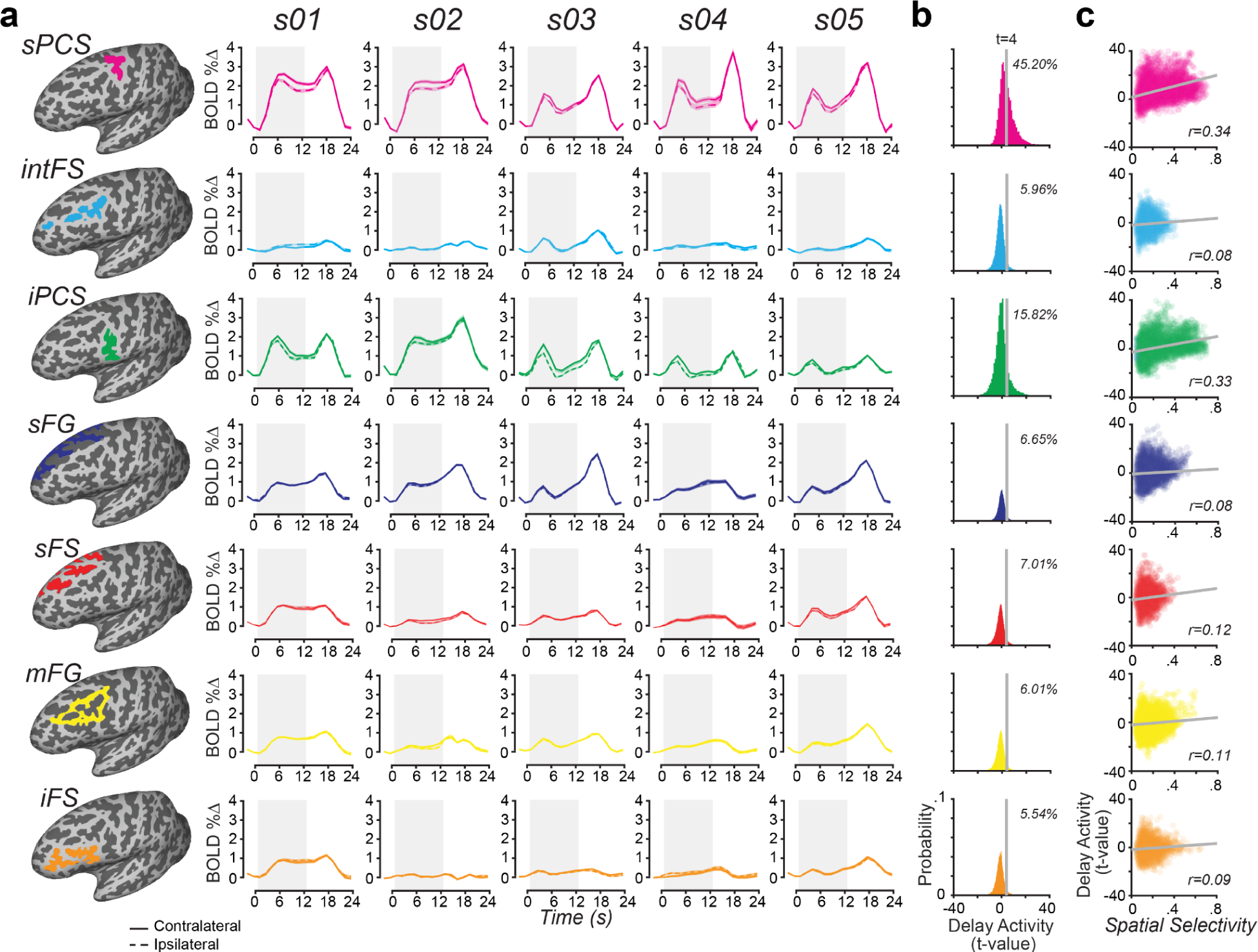
Delay period activity in all ROIs in all participants, related to Figure 3. (a) Trial-averaged (± s.e.m.) BOLD time courses from all ROIs, across five participants, time-locked to trial onset. Solid and dashed lines correspond to trials with targets in the contralateral and ipsilateral visual fields, respectively. Error bars indicate s.e.m across trials. Gray shaded box denotes the delay period. (b) Histograms of t-values of delay activity pooled across participants. Gray line represents the threshold and insets are the percentage of voxels which are above the threshold (c) Delay activity plotted against spatial selectivity. Pearson’s r was calculated between t-values of delay activity and variance explained in the pRF model. Gray line represents the best linear fit.

**Supplementary Figure 4.**
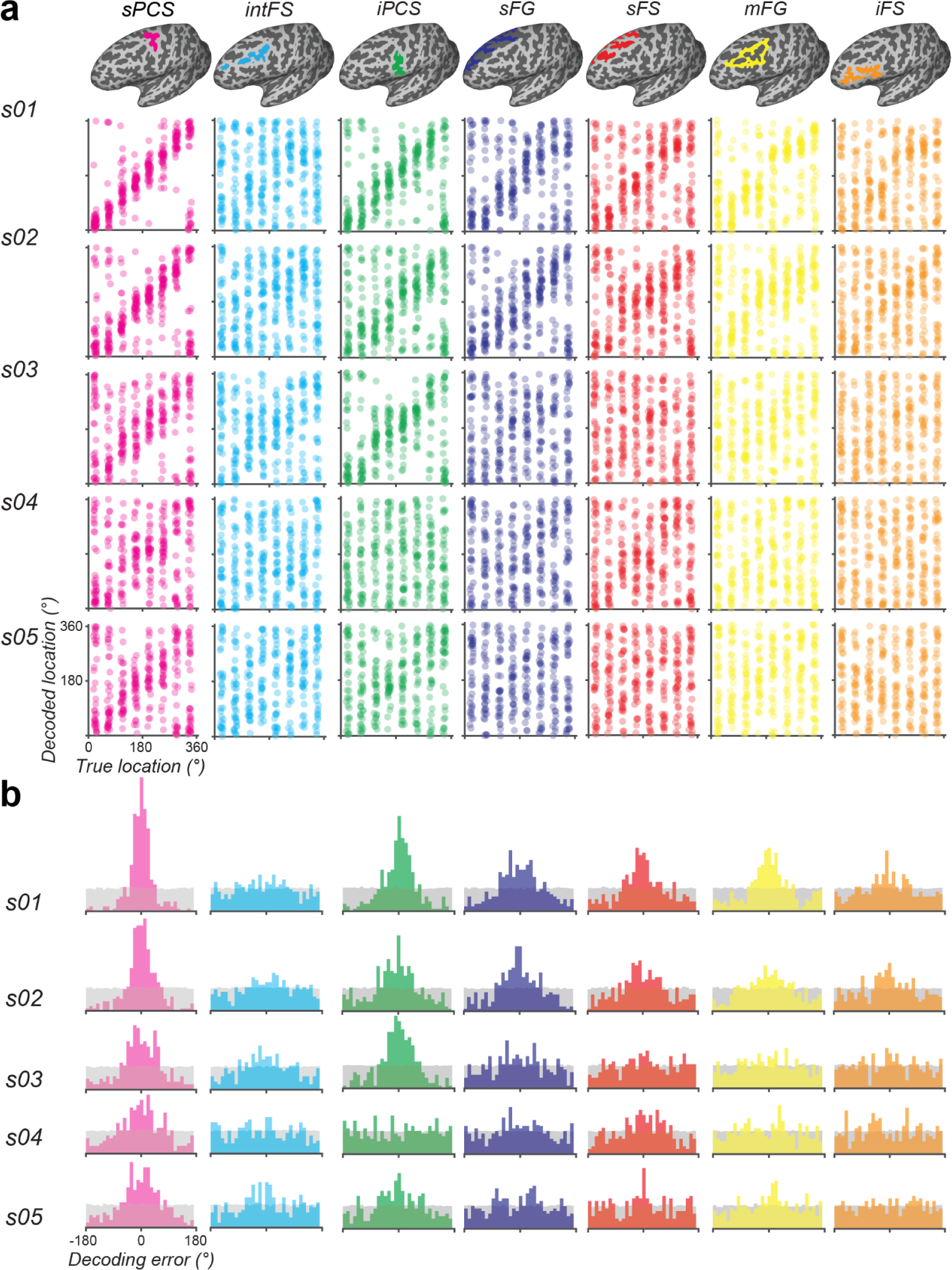
Decoding performance in all ROIs, related to Figure 4 a & b. (a) Decoded target locations plotted against true target locations for each participant. (b) Distribution of decoding errors compared to a null distribution from label-shuffled data (gray).

**Supplementary Figure 5.**
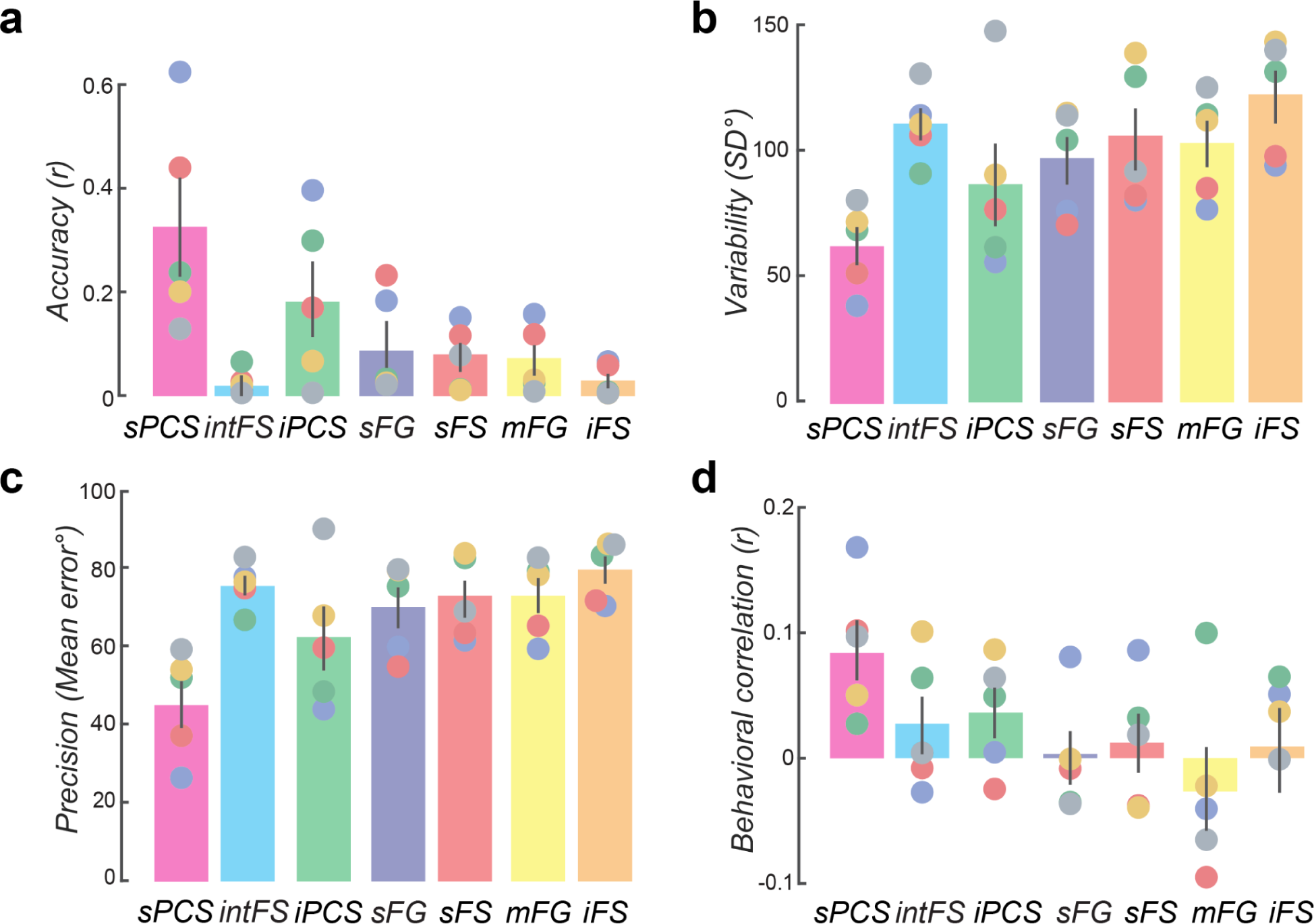
Decoding performance in all ROIs, related to Figure 4 c - f. (a) decoding accuracy (circular correlation between decoded and true location), (b) decoding precision (mean absolute error), and (c) decoding variability (standard deviation of signed decoding errors). Error bars indicate ± sem. (d) Circular correlation between the behavioral memory errors and neural decoding errors across participants. Colored dots represent individual participants; gray bars represent group means; error bars indicate ± sem.

**Supplementary Figure 6.**
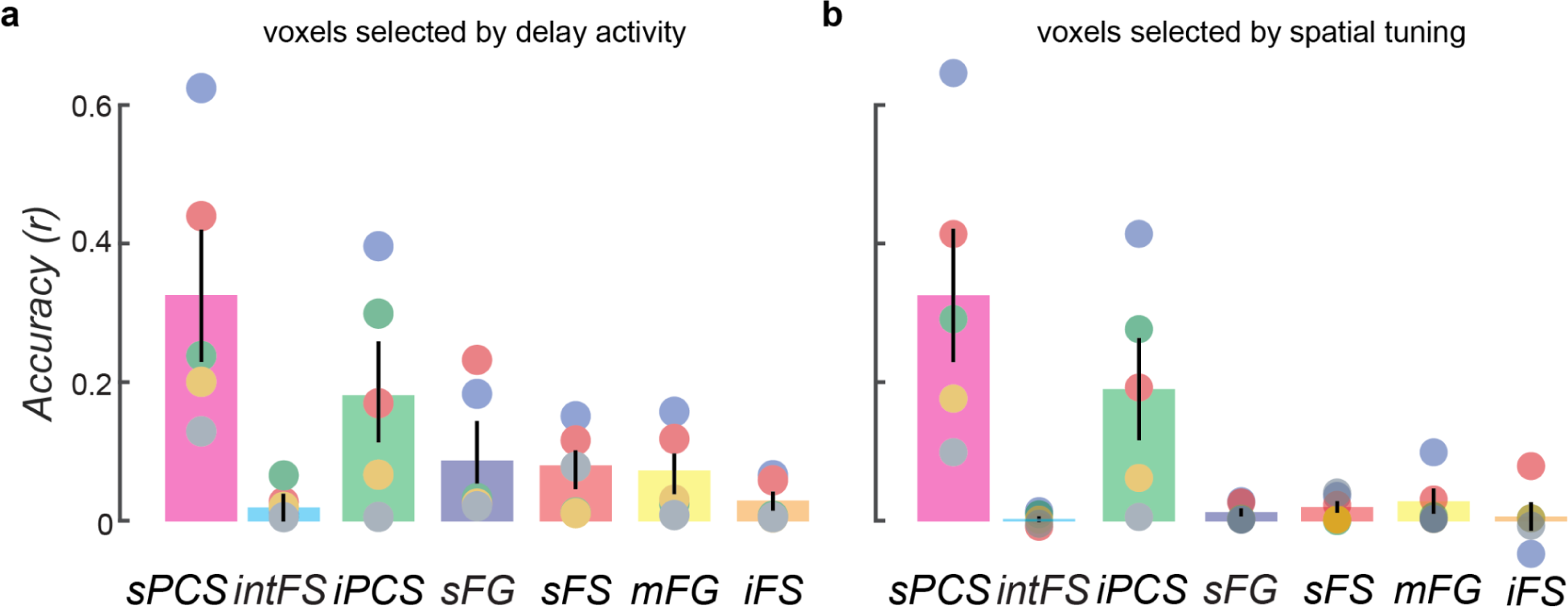
Decoding performance in all ROIs by different voxel selection strategies, related to Figure 4c. (a) Top 1000 voxels that exhibited the strongest delay-period responses, as determined by their t-values from the GLM contrast comparing delay-period activity to baseline. (b) Top 1000 voxels whose pRF model fit surpassed a common threshold (pRF R² ≥ 10%).

**Supplementary Figure 7.**
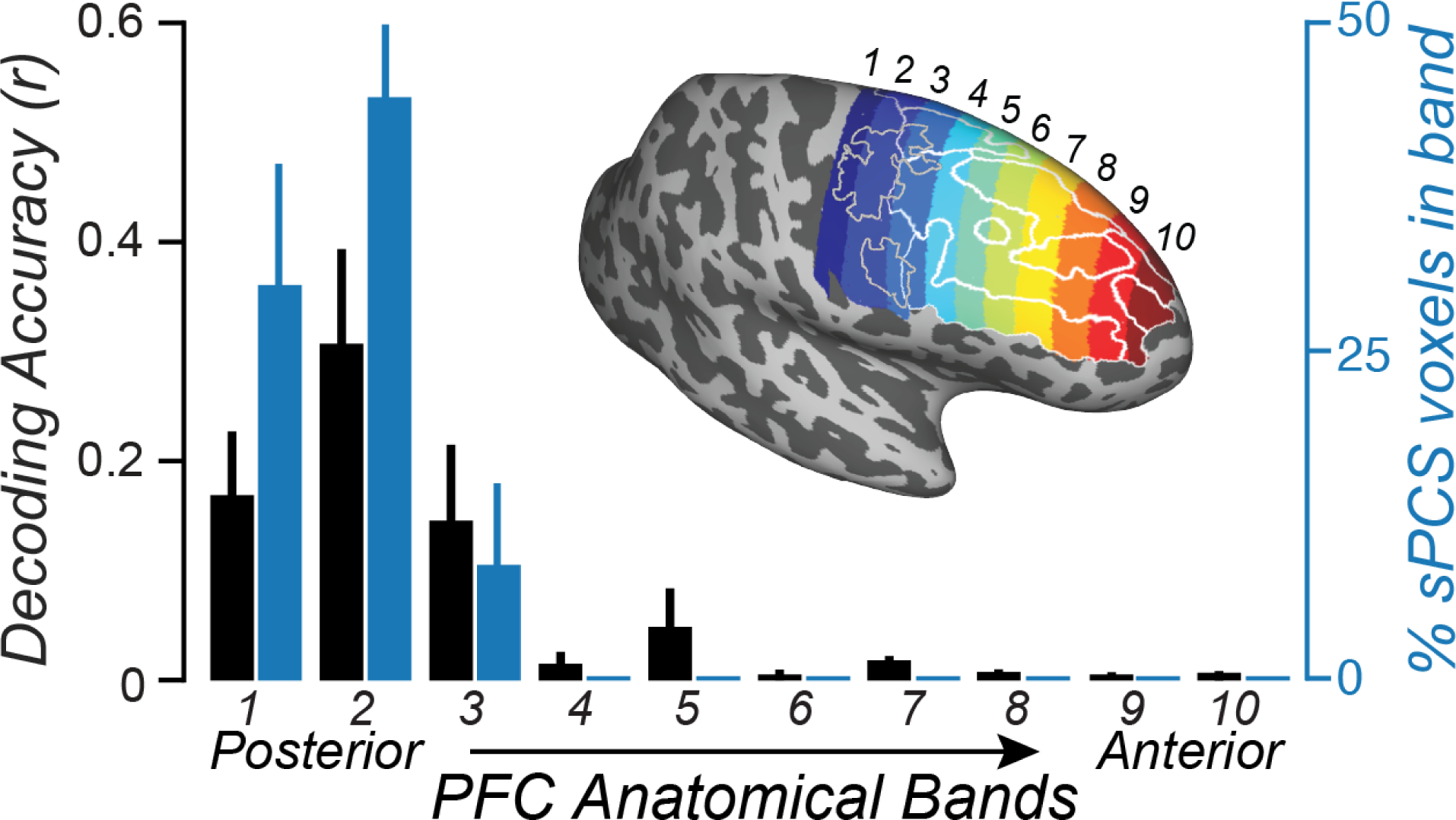
Decoding performance across anatomical bands in PFC. The PFC was segmented into ten 22 mm wide bands. The brain inset depicts the numbered color-coded bands as well as ROI boundaries of the anatomical regions (white lines) and pRF defined visual field maps in the precentral sulcus (gray lines) for a participant s01. Decoding was performed within each band and without regard for the ROIs. The black bars are decoding accuracy (circular correlation between decoded and true location). Note that decoding was only successful in posterior bands 1, 2, and 3 (p=0.001, p=0.001, and p=0.003 respectively). The blue bars are the percentage of voxels in each band that belong to the sPCS visual field map. The bars represent group means; the error bars are ±SEM. Notice that decoding success depends on the percentage of sPCS voxels in the band.

**Supplementary Figure 8.**
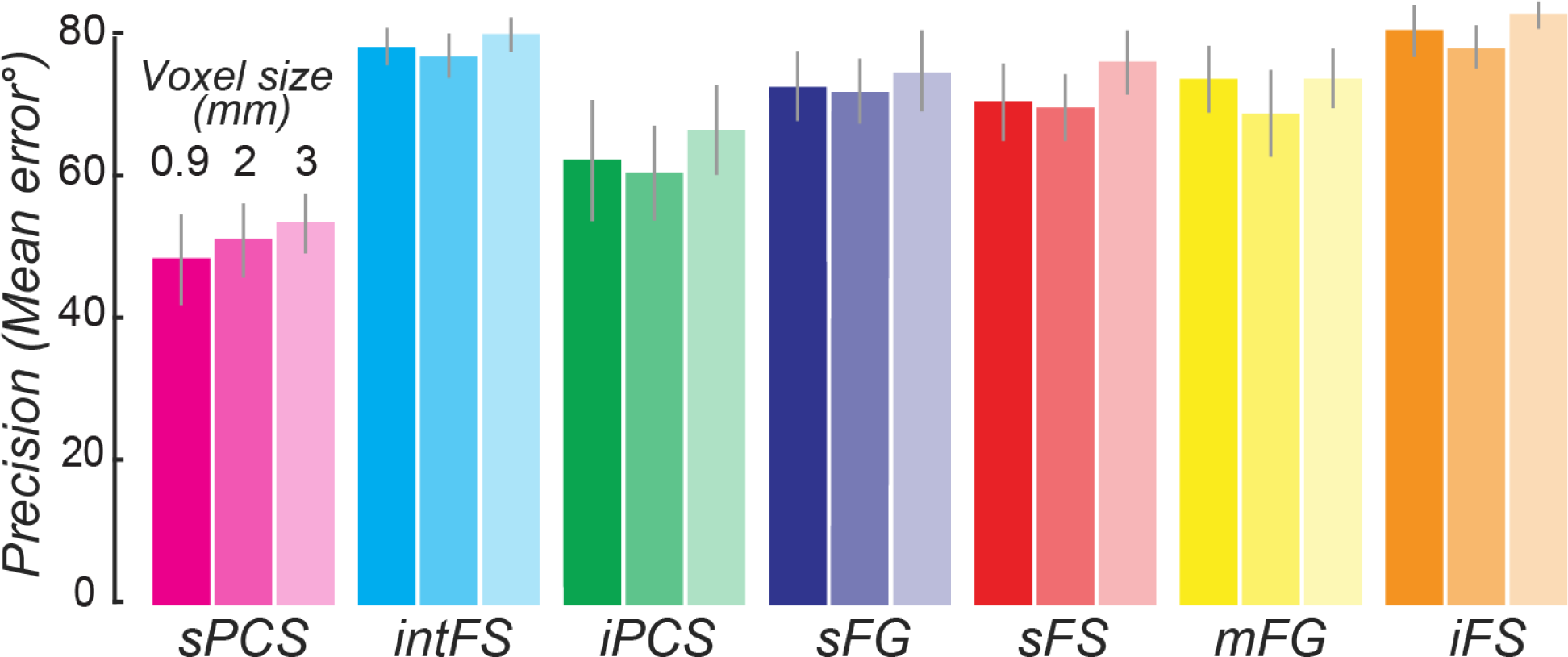
Decoding precision at resampled voxel size 2 and 3 mm, related to Figure 4. Decoding procedure and precision were the same as in Figure 4.

**Supplementary Figure 9.**
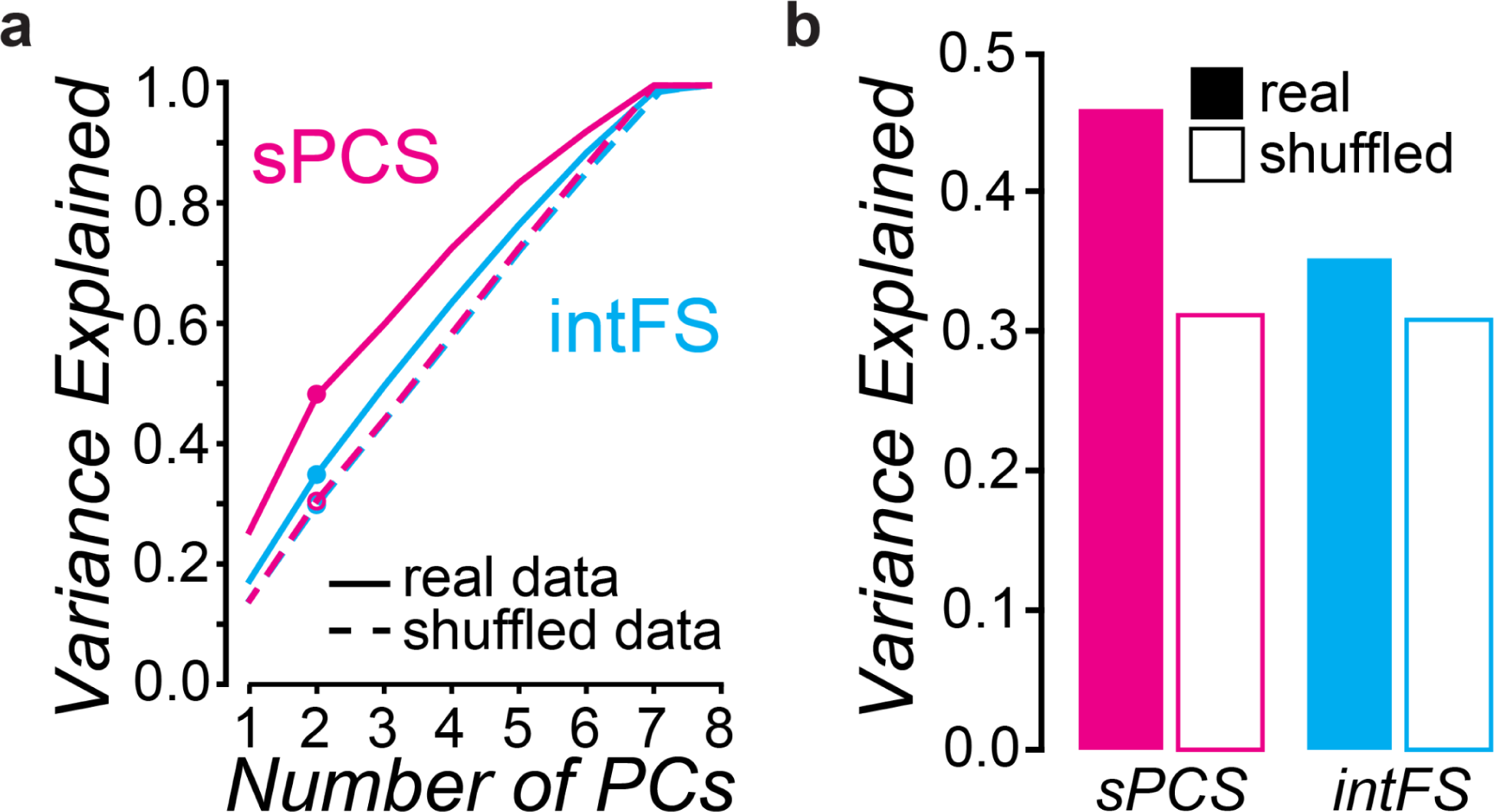
Principal component analysis, related to Figure 5 b & d. (a) Scree plots in sPCS and intFS. (b) Variance explained by the first two PCs.

**Supplementary Figure 10.**
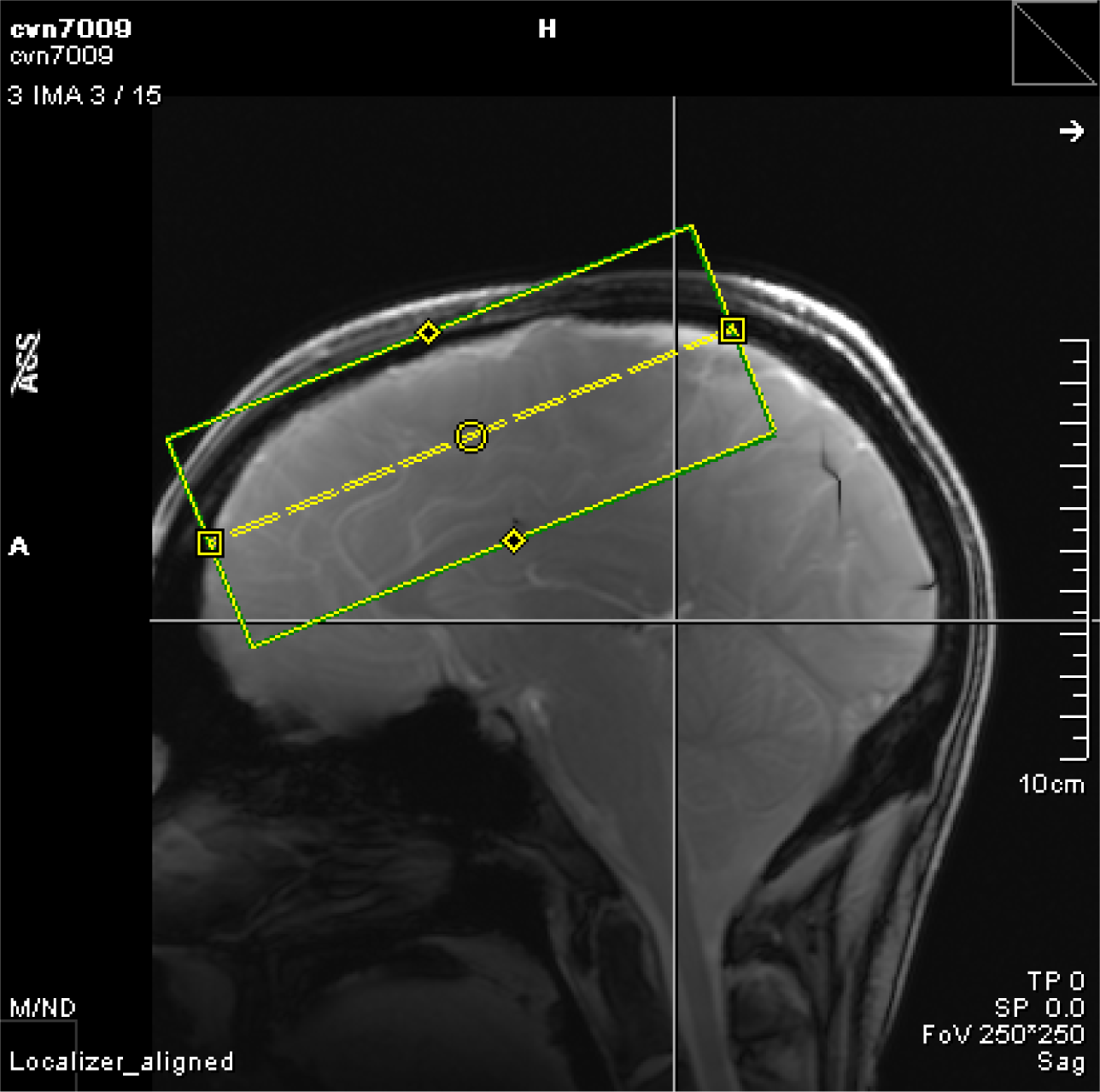
Field-of-view for the fMRI data acquisition at 0.9mm. The green box delineates the slab of prefrontal cortex in a representative participant that we were able to image at high resolution (0.9mm isotropic voxels) and reasonable temporal resolution (2100ms TR). The partial brain coverage was composed of 60 slices, with a slice thickness of 0.9mm, slice gap 0mm, and a field-of-view 135mm (FE) x 133.2mm (PE).

## Notes

### Competing Interest Statement

The authors have declared no competing interest.

